# Large-scale analysis of optimisation methods for parameter estimation problems in the life sciences

**DOI:** 10.64898/2026.07.11.737731

**Authors:** Stephan Grein, David R. Penas, Daniel Weindl, Polina Lakrisenko, Julio R. Banga, Jan Hasenauer

## Abstract

Dynamic models are central to the computational life sciences but typically contain unknown parameters that must be inferred from experimental data. High-throughput measurements have made this task increasingly challenging, yielding high-dimensional search spaces and non-convex objectives with many local optima. This makes the choice of optimisation method critical. However, existing empirical studies either consider only a limited number of benchmark problems or only a narrow spectrum of local, global and hybrid optimisation methods.

Here, we present a comprehensive benchmark of a broad range of optimisation methods on a curated collection of parameter estimation problems, comprising 990 method-problem-pairs executed on two independent supercomputing infrastructures. Our evaluation quantifies success rates, solution quality and computational cost, revealing characteristic strengths and limitations of each approach. We find that optimisation methods separated into clear performance tiers. Building on these results, we implemented a new hybrid strategy that combines enhanced scatter search with the best-performing local solver, which showed robust performance and improved on the other scatter-search variants we tested. Our results provide practical guidance for selecting optimisation methods and thereby support more accurate and reliable model calibration.

## Introduction

Systems biology, systems medicine and epidemiology are interdisciplinary fields that aim to understand complex processes using mathematical modelling Qiao et al. (2025). The spectrum of modelling approaches is broad and includes ordinary differential equations (ODEs), partial differential equations (PDEs), stochastic models and agent-based models. These models typically possess parameters that cannot be measured directly, and therefore, need to be estimated to accurately represent the underlying systems. Accordingly, a central task in the model development process is parameter estimation (Tarantola, 2005), where the goal is to determine parameter values that best reproduce observed data. Accurate parameter estimation is crucial for model validation, prediction accuracy and the design of biological experiments (Qiao et al., 2025; Lopatkin and Collins, 2020).

Mathematical models and parameter estimation problems have steadily grown more challenging due to the advent of novel high-throughput measurement techniques (Goodwin et al., 2016; Karahalil, 2016). Nowadays, parameter estimation problems are often high-dimensional (see, e.g. Bouhaddou et al. (2018); Fröhlich et al. (2023); Berndt et al. (2018)) and frequently give rise to non-convex objective functions with multiple local optima (Moles et al., 2003; Hass et al., 2019), which complicates the optimisation process. To address these challenges, a broad spectrum of optimisation methods has been proposed (see review by Fröhlich et al. (2019)). These optimisation methods can broadly be categorised into local, global and hybrid techniques.

Local optimisation algorithms, such as the Levenberg–Marquardt algorithm (Levenberg, 1944; Marquardt, 1963) and trust-region methods (Conn et al., 2000), are efficient and effective when the objective function is smooth and the initial parameter guess is close to the global optimum. These methods typically rely on gradient information to navigate the search space and can achieve rapid convergence. However, they are prone to getting trapped in local minima, especially in the complex landscapes that are typical for parameter estimation problems in systems biology. As a result, standard naïve local optimisation algorithms often fail to find the global optimum. A simple yet effective strategy to enhance local optimisation is the use of multi-start methods (Raue et al., 2013). In this approach, local optimisation is performed from multiple, randomly chosen starting points in parameter space (Ullah and Seidel-Morgenstern, 2009), for example, sampled from a Latin hypercube (Mckay et al., 1979). This increases the likelihood of finding a global solution. While multi-start methods can substantially improve the robustness of local optimisation techniques, they can be computationally expensive, particularly for high-dimensional problems.

Global optimisation methods are designed to explore the entire search space without getting trapped in local minima and to locate the global optimum. Techniques such as evolutionary programming (Fogel et al., 1991), genetic algorithms (Holland, 1975), particle swarm optimisation (Kennedy and Eberhart, 1995), simulated annealing (Kirkpatrick et al., 1983), and scatter search (Egea et al., 2007) have been employed successfully in this context. These metaheuristics do not require gradient information and are well suited for handling multi-modal objective functions, but they often come with higher computational costs.

Hybrid optimisation methods seek to combine the efficiency of local and the reliability of global optimisation techniques. Their development has gained traction in particular for dynamical systems. It has been shown that combining global search strategies with local refinement steps can offer a favourable trade-off between exploration and exploitation (Villaverde et al., 2018). For deployment on high-performance computing (HPC) infrastructure, methods that support asynchronous communication have also been developed (Penas et al., 2017).

Despite this wealth of methods, researchers are often left without clear guidance on method selection. Rigorous head-to-head comparisons remain scarce. Quantitative benchmarking is essential to assess performance and guide method selection in systems biology. Early efforts introduced benchmark collections but covered only a narrow slice of the algorithmic landscape: Hass et al. (2019) evaluated two local optimisation methods on 20 benchmark problems, while Villaverde et al. (2018) compared six optimisation methods (with two different gradient computation schemes) on seven benchmark problems. Community-curated benchmark suites reflecting published estimation problems have since been assembled and standardised (Schmiester et al., 2021), facilitating fairer evaluations. Nevertheless, gaps persist: prior studies explored only restricted configurations and largely overlooked parallelisation, which is increasingly critical as model and dataset sizes grow.

In this study, we provide a comprehensive evaluation of a broad spectrum of parallel optimisation methods suitable for high-performance computing infrastructures. To assess the reliability and efficiency of these methods, we considered 30 published benchmark problems for ODE-constrained optimisation problems. These benchmark problems originate from systems biology, systems medicine and systems epidemiology and exhibit a broad range of characteristics. Our results elucidate the strengths and weaknesses of each method and offer guidelines for their application in different scenarios. This not only saves computational resources but also accelerates the iterative cycle of model development and validation.

## Results

### Pipeline for optimiser benchmarking

To perform a comprehensive evaluation of optimisation methods for dynamical modelling in systems biology, systems medicine, and epidemiology, we implemented a computational pipeline that can be deployed on different computing infrastructures with minimal adaptation. The pipeline builds on the Python Parameter Estimation Toolbox `pyPESTO` (Schälte et al., 2023) and the Parallel Parameter Estimation package `parPE` (Schmiester et al., 2020), which in turn use the Advanced Multi-language Interface `AMICI` (Fröhlich et al., 2021) for the `CVODES` solver for stiff and non-stiff ODE systems (Serban and Hindmarsh, 2005).

The pipeline facilitates the streamlined evaluation of a broad spectrum of state-of-the-art optimisation algorithms. Via `pyPESTO` it provides access to the `NLopt` library for nonlinear optimisation (Johnson, 2023) and the `SciPy` optimisation algorithms (Virtanen et al., 2020). These free, open-source packages offer a comprehensive collection of local and global optimisation algorithms. We complemented this collection with optimisation algorithms that are commonly used in, or even have been designed for, applications in systems biology. This includes the covariance matrix adaptation evolution strategy `CMA-ES` (Hansen et al., 2019), the particle swarm algorithm `PySwarm` (Lee, 2013), the interior point algorithm `Ipopt` (Wächter and Biegler, 2006), and the trust-region algorithm `fides` (Fröhlich and Sorger, 2022), which are interfaced via `pyPESTO`, and the parallelised enhanced scatter search algorithm `SaCeSS` (self-adaptive cooperative enhanced scatter search, Penas et al. (2017)) with different local optimisation algorithms that uses `parPE`.

In this study, we consider 33 optimisation methods that differ in both the underlying optimisation algorithm and the parallelisation scheme (Figure 1A). Most optimisation methods were implemented as parallel multi-start optimisations via `pyPESTO`, distributing independently initialised optimiser starts across cores, whereas the three `SaCeSS` variants natively exploit MPI-based parallelism through coordinated search processes (Figure 1C,D). In the multi-start setup, individual optimisation starts were distributed across cores and executed sequentially on each core; objective function evaluations themselves were not parallelised. The majority of optimisation algorithms were provided by `NLopt` (n=15) and `SciPy` (n=11) (Figure 1B). Overall, the considered optimisation strategies comprised local (n=28), global (n=3), and hybrid (n=2) approaches. Most local optimisation algorithms used first-order or first- and second-order derivative information (n=19).

**Fig 1:**
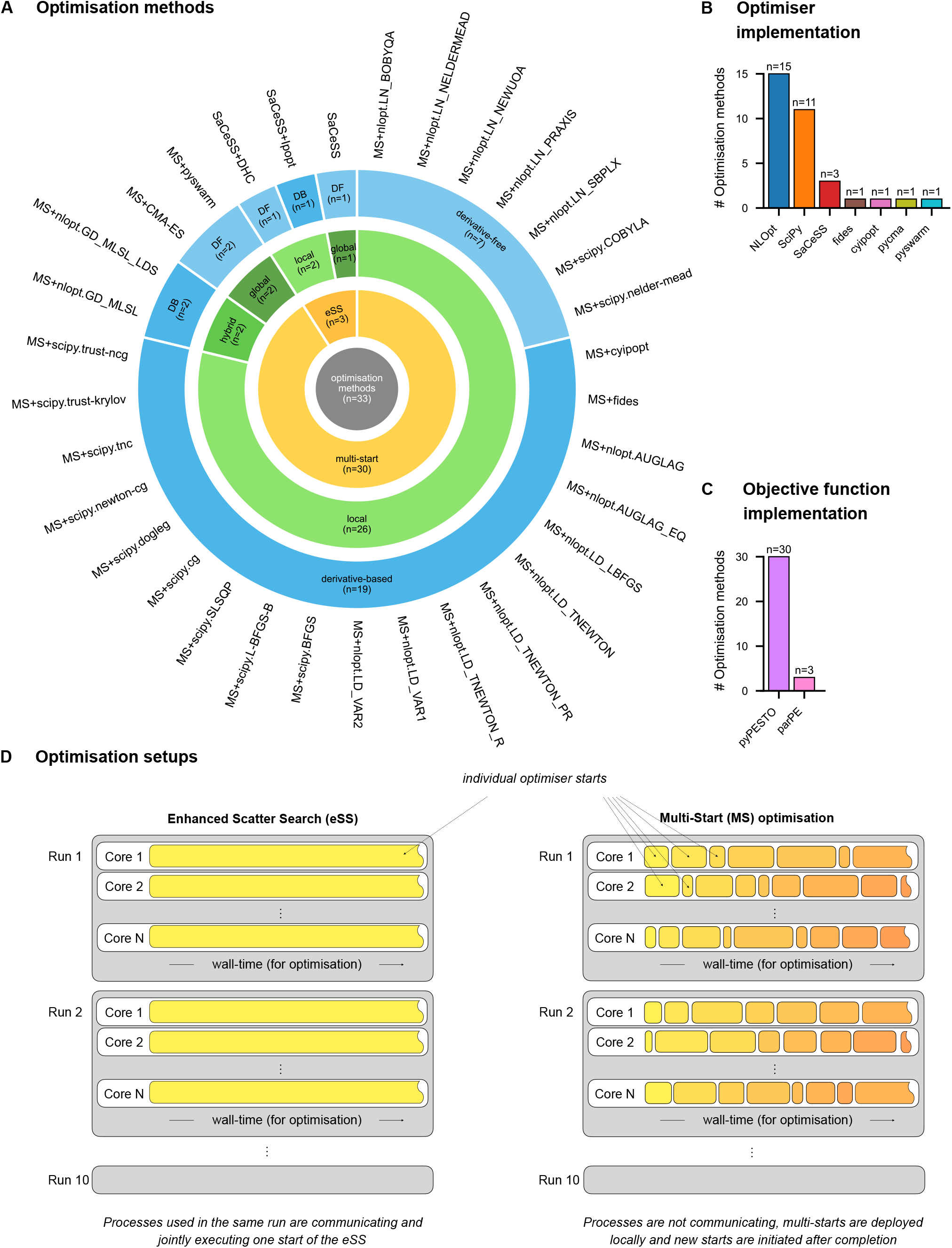
Optimisation methods considered in this study. (A) Optimisation methods considered in this study and their properties (DB: derivative-based, DF: derivative-free). The MS+ prefix indicates multi-start methods. `nlopt.LD_TNEWTON_{RESTART,PRECOND_RESTART}` are abbreviated as `nlopt.LD_TNEWTON_{R,PR}` throughout the manuscript. (B) Number of optimisation algorithms from different optimisation libraries considered in this study. (C) Number of optimisation methods building on objective function implementations in `pyPESTO` or `parPE`. (D) Illustration of setups for parallel, enhanced scatter search (eSS) (as implemented in `SaCeSS`) and multi-start optimisation. A single eSS run exploits all available cores and communication between them to execute a single start until the wall time limit is reached, while a multi-start run uses a large number of starts across cores and time points.

For a realistic assessment of the optimisation methods, the pipeline uses the PEtab benchmark collection of estimation problems in systems biology, systems medicine, and epidemiology (Figure 2A). It uses the parameter estimation table (PEtab) format (Schmiester et al., 2021) for a standardised encoding of the estimation problem. The PEtab specification includes model, parameter bounds and scales, experimental conditions, and measurement data. Here, we consider 30 parameter estimation problems contained in the PEtab benchmark collection. The considered benchmark problems contain 3–155 unknown parameters which are subject to optimisation and differ also with respect to the number of state variables, observables, experimental conditions, and data points (Figure 2B). Furthermore, they include known and parameter-dependent initial conditions (including pre-equilibration), as well as different noise models (Figure 2C). This collection covers a broad spectrum of problem characteristics, and captures expected correlation patterns, e.g., between the number of parameters and measurements (Supplementary Figure S1). The objective functions for the optimisation are either negative log-likelihood or negative (non-normalized) log-posterior probability, depending on the availability of prior information on the parameters.

**Fig 2:**
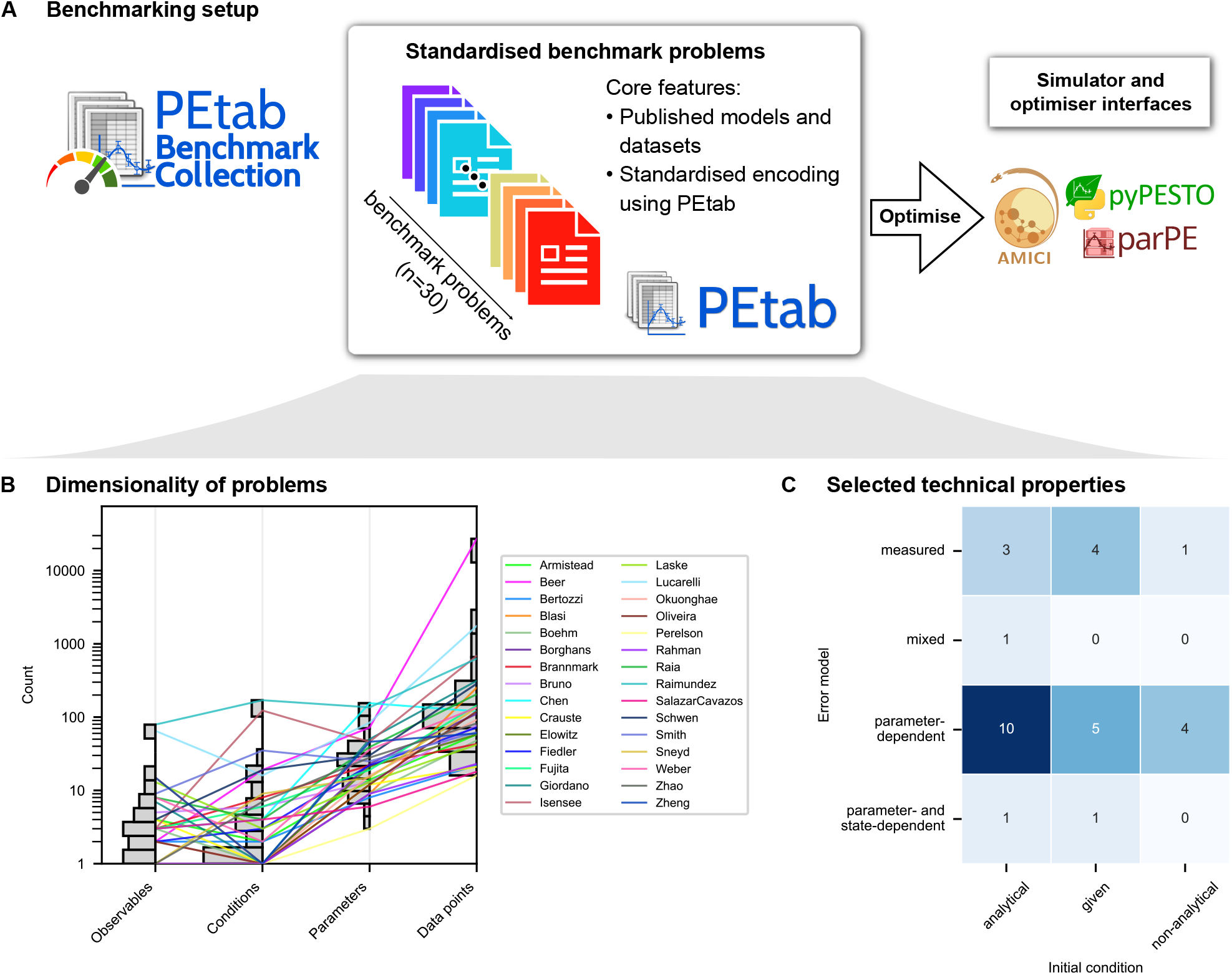
Benchmark problems considered in this study. (A) Source and encoding of benchmark problems, as well as software tools used for model simulation and objective function (gradient) evaluation. (B) Dimensionality of the benchmark problems. (C) Technical properties of the benchmark problems, i.e., type of error model and initial conditions.

In summary, the implemented pipeline facilitates the evaluation of a comprehensive collection of optimisation methods across a broad spectrum of application-derived benchmark problems. The use of state-of-the-art numerical methods for model simulation, objective function evaluation and gradient computation ensures that the benchmark results are relevant for current parameter estimation workflows in systems biology, systems medicine and epidemiology. In the following, we distinguish between optimisation algorithms, referring to the underlying numerical methods (e.g., `scipy.BFGS`, `Ipopt`, `fides`), and optimisation methods, referring to the full computational setups in which these algorithms are deployed (e.g., `MS+scipy.BFGS`, `MS+Ipopt`, `SaCeSS+Ipopt`).

### Benchmark design and computational budget allocation

To assess the performance of the optimisation methods for the collection of benchmark problems, we employed the afore-described pipeline to study all 990 combinations of 30 benchmark problems and 33 optimisation methods. For the selection of plausible compute time budgets, we assessed the computation time for the simulation of the models as well as the computation of the objective functions and the objective function gradients. Furthermore, we collected information about the expected problem complexity from the original publications.

The assessment of the computation times revealed that model simulations required between 0.047 ms and 769 ms per experimental condition (Supplementary Figure S2A). As the objective function evaluation requires the simulation of multiple experimental conditions, the computation time for an objective function evaluation ranged from 0.253 ms to 2203 ms (Supplementary Figure S2B). The range widened as the parameters of large models are usually inferred from more experimental conditions (Supplementary Figure S2C,D). The computation time for gradient evaluation was determined for forward and adjoint sensitivities (Supplementary Figure S3A,B). Interestingly, we did not only find a difference in computation time which strongly depended on the number of parameters (Supplementary Figure S3D), but for some problems, gradient calculation using adjoint sensitivities (mean success rate 86%) proved more reliable than with forward sensitivities (mean success rate 74%) (Supplementary Figure S3B). Based on these results, we selected for each benchmark problem the more efficient and more reliable gradient computation method, yielding computation times of 0.723 ms to 491 s for a joint objective function and gradient evaluation. Based on this and the results of previous studies, the benchmark problems were divided into two groups – simple problems (*n* = 23) and challenging problems (*n* = 7) – each allocated a different computational budget: simple problems were run on 12 cores with a wall-time limit of 3 hours, while challenging problems were run on 24 cores with a wall-time limit of 9 hours.

Given the sensitivity analysis methods and compute budget, we performed a comprehensive benchmarking of the 990 combinations of benchmark problems and optimisation methods. The computations were performed on the HPC clusters *Marvin* of the University of Bonn (Germany), and *FinisTerrae III* of the Galicia Supercomputing Center (CESGA, Spain). The analysis was repeated 10 times with randomly assigned random number generator seeds, yielding 10 runs which could be used to assess the stochasticity of the outcome. Overall, this benchmarking required over 1.5 million core-hours and produced approximately 30 TB of raw data files, and would not have been feasible without the utilization of HPC infrastructure. These result files provide detailed information about the optimiser trajectories, including parameter vectors and objective function values for different multi-starts, iterations and time points. These results were filtered, to remove infeasible solutions which have been reached due to an optimisation algorithm not respecting parameter constraints or wall time limits. In the following, we present an in-depth assessment of these data to assess the suitability of different optimisation methods for problems in systems biology.

### Reproducibility of optimisation results

Reproducibility is a cornerstone of scientific inference (Antunes and Hill, 2024), yet, even for the same optimisation methods, reported performances vary widely across studies. Such discrepancies can stem from differences in study design, implementation details, and computing environments. In computing, sources of variability range from hardware and software versioning to non-associative floating-point operations and out-of-order arithmetic (Goldberg, 1991). To ensure the robustness of our findings, we evaluated the reproducibility of optimisation performance across two independent HPC systems, *Marvin* and *FinisTerrae III*, and compared the resulting outcomes (Figure 3A).

**Fig 3:**
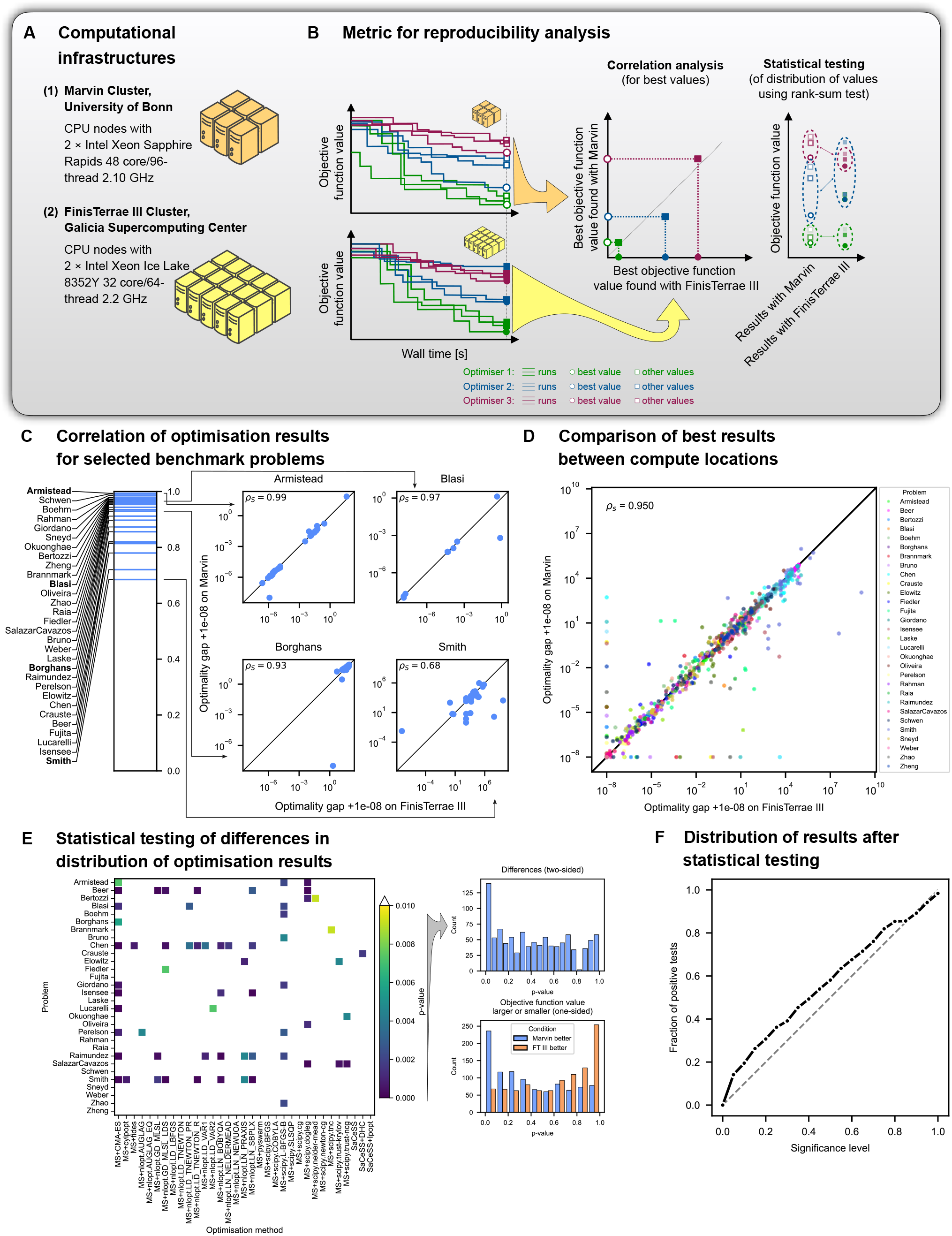
Reproducibility analysis. (A) Information about the computing clusters used for this study. (B) Illustration of recorded data for optimisation runs and metric to assess reproducibility across HPC systems. (C) Spectrum of observed Spearman correlation coefficients of the minimal optimality gaps of optimisation methods across HPC systems, and four exemplary scatter plots showing the underlying data. (D) Agreement of best results for all benchmark problems across compute sites. (E) Assessment of differences between the distribution of optimality gaps obtained from the two HPC systems: (left) Heatmap and (right) histograms. P-values were calculated using the two-sample, one- and two-sided rank sum test. (F) Comparison of significance level and fraction of positive test for two-sample, two-sided rank sum test results in (E, right). The reference points for the optimality gaps were the best values achieved by any run of the optimisation methods shown in (E) on either computing cluster.

To evaluate the reproducibility, we computed the correlations and performed statistical tests (Figure 3B). Among other things, we studied the optimality gap of the *i*-th run of the *j*-th optimisation algorithm at the *k*-th location (HPC systems),

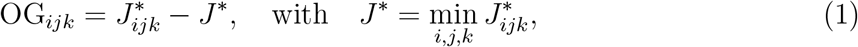

which measures the difference between the objective value found by the optimisation run, 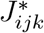, and the best objective value found for this problem, *J*^*∗*^. The optimality gap OG_*ijk*_ is zero if the best objective function value is reached, and otherwise positive. Note that we did not use the distance of the parameter vectors as a measure as several benchmark problems are known to be subject to non-identifiability (Kreutz, 2018), meaning that the optimal parameter vectors are not unique and non-zero distances are to be expected and not indicative of non-convergence per se.

Our analysis of the benchmarking results indicated a good agreement between the two HPC systems. The minimal optimality gaps for algorithms across the 10 runs, 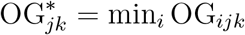, were highly correlated between the two systems. For the individual benchmark problems, we observed a Spearman correlation between 0.685 and 0.994 (Figure 3C), with a median of 0.955. For the full collection of benchmark problems, we found a Spearman correlation of 0.95 (Figure 3D).

For an assessment of the results beyond the best run, we studied the distribution of optimality gaps across the 10 runs. As the runs used randomly sampled starting values, the observed _10 10_ optimality gaps – 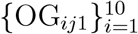 for *Marvin* and 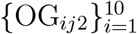 for *FinisTerrae III* – should follow the same distribution if the execution is reproducible. Statistical testing using the two-sample and two-sided rank sum test revealed a small number of significant differences (Figure 3E, left). For a significance level of *α* = 0.01, we found 62 combinations of benchmark problems and optimisation methods for which the differences in the distribution of optimality gaps were statistically significant. This corresponds to 6.3 % of the combinations, and is therefore higher than expected. The p-value distribution shows some enrichment close to 0 and 1 (Figure 3E, right, top). This disagreement with the uniform distribution is increasing when considering the corresponding one-sided tests (Figure 3E, right, bottom). It appears that each infrastructure slightly favours a subset of the problems (without a clear tendency). Yet, the assessment of the fraction of positive tests for different significance levels (Figure 3F) still shows a good agreement.

In summary, we found good agreement of the optimisation results at *Marvin* and *FinisTerrae III*. Although the runs do not seem to be sampled from the exact same distribution, the minor difference can easily be explained by the difference in the performance parameters of the employed supercomputers. Due to the good agreement also found for subsequent analyses, we present in the main manuscript only the results for *Marvin*, and provide additional results for *FinisTerrae III* as Supplementary Data.

### Optimality of optimisation results

Optimality of optimisation results is crucial for the assessment of a model’s ability to describe the experimental data, and therefore, for tasks such as model selection and falsification. In fact, as the limitations of models and optimisation method cannot be easily distinguished in applications, suboptimal optimisation results can cause the rejection of valid models. Here, we assessed the ability of optimisation methods to recover the optimal solution of the parameter estimation problem using the minimal optimality gap across the 10 runs, 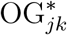 (Figure 4A). A non-zero optimality gap indicates that the achieved solution is suboptimal.

**Fig 4:**
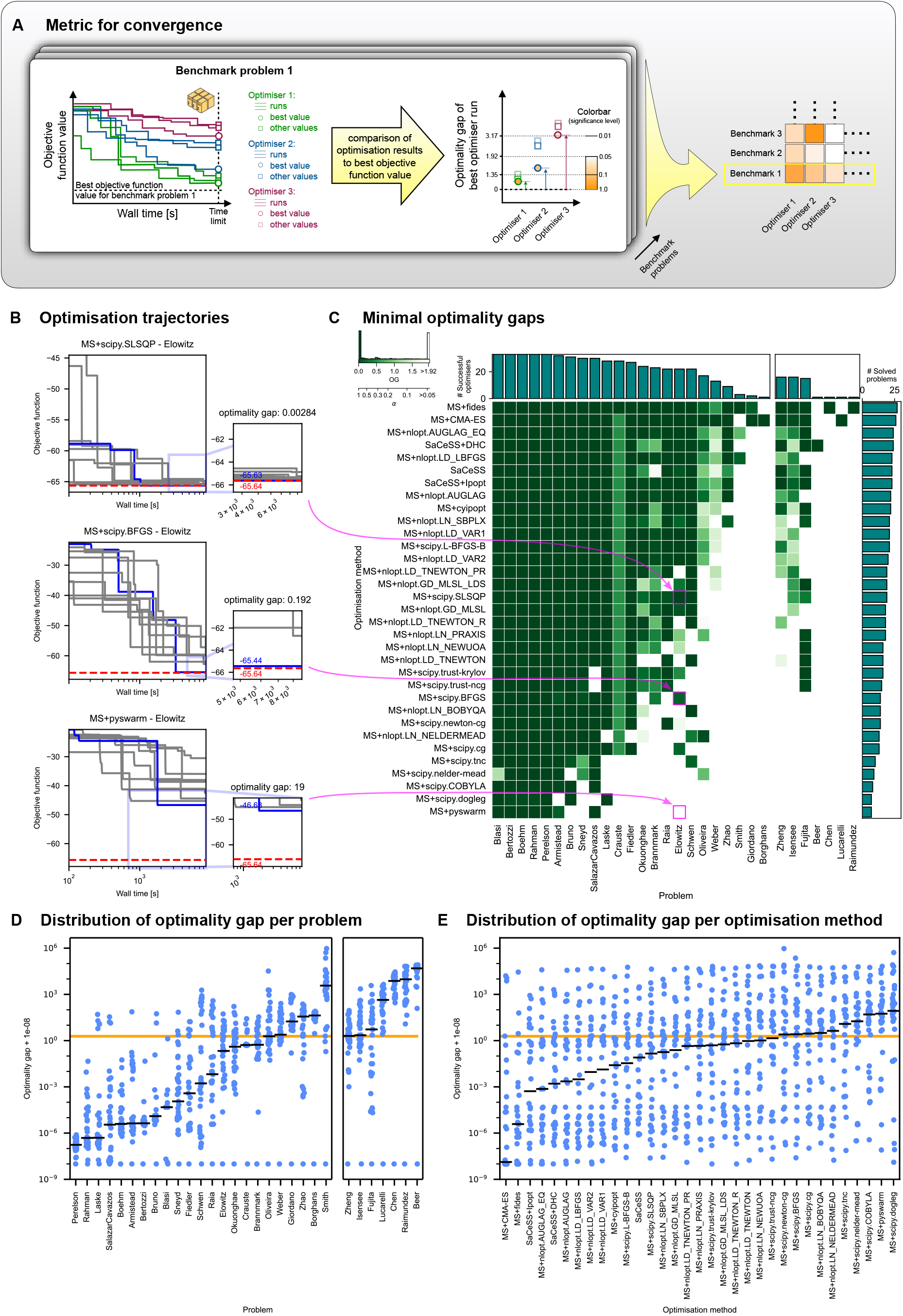
Optimality analysis. (A) Illustration of optimisation runs and minimal optimality gap. (B) Convergence curves for three combinations of optimisation method and benchmark problem. The combinations differ with respect to the minimal optimality gap and number of successful runs. The solid lines represent the trajectories of the 10 runs. The optimality gap is the distance between the red line (best objective value found for the given problem across optimisation methods) and the end point of the blue line (best value from the current optimisation method). (C) Heatmap of the minimal optimality gap across 10 runs for all combinations of benchmark problems and optimisation methods; bar plots of the number of successful optimisation methods per problem and number of solved problems per optimisation method. A problem was considered solved if the minimal optimality gap across the 10 runs was below 1.92, which corresponds to a *χ*^2^-based rejection test with significance level 0.05. (D) For each problem, the minimal optimality gap found across the 10 runs of each optimisation method (*n* = 33 each). (E) For each optimisation method, the minimal optimality gap found across the 10 runs for each problem (*n* = 30 each). The orange lines in (D) and (E) indicate the optimality gap under which a problem was considered solved. The reference points for the optimality gaps were the best values achieved by any run of the optimisation methods shown in (E) on Marvin.

Our analysis of the benchmarking results revealed substantial differences between optimisation methods with respect to convergence curves and optimality gaps (Figure 4B). Even for a significance level of 0.05 in a *χ*^2^-based rejection test (Villaverde et al. (2018)), corresponding to an acceptable optimality gap of 1.92, we found that the 33 optimisation methods solved on average 17.6 out of the 30 benchmark problems (range: 7 to 27) (Figure 4C), corresponding to 59 % (range: 23% to 90%). The top performing optimisation methods with respect to the number of successfully solved problems were `MS+fides` (*n* = 27), `MS+CMA-ES` (*n* = 26), `SaCeSS+DHC` (*n* = 24), `MS+nlopt.AUGLAG_EQ` (*n* = 24), `SaCeSS` (*n* = 23), `SaCeSS+Ipopt` (*n* = 23), and `MS+nlopt.LD_LBFGS` (*n* = 23). The top performers cover a broad range of different methods, including enhanced scatter search and multi-start methods using gradient-free and gradient-based optimisation algorithms. Interestingly, various widely used methods solved only a relatively low number of problems successfully, e.g., `MS+pyswarm`. As expected, we also observed a substantial difference between the benchmark problems. The benchmark problems with the lowest number of successful optimisation methods (≤ 2), namely Beer et al., Borghans et al., Chen et al., Lucarelli et al., Raimundez et al., Giordano et al., were only solved by `MS+CMA-ES`, `SaCeSS+DHC` or `MS+fides`. For these problems, there was a substantial degree of variability between runs as well as between HPC systems (see comparison of Figure 4 and Figure 3D). Importantly, optimisation methods that solved the challenging problems were usually also successful on the simple problems, but not vice versa; the sets of solved problems were largely nested, not methods-specific. A fuzzy ranking of optimisation methods solely on the optimality gap confirmed that the ranking by the number of solved problems was not dominated by small variations around the chosen optimality gap cut-off (Supplementary Figure S4). Indeed, the ranking was quite stable over a wide range of optimality gap thresholds (Supplementary Figure S6).

To assess the variability across runs, we complemented the study of the minimal optimality gaps with an analysis of the distribution of optimality gaps achieved on *Marvin* and *FinisTerrae III*. Interestingly, even for benchmark problems that were successfully solved by most optimisation methods, when looking at the best run, we have a broad distribution of optimality gaps (Figure 4D), e.g., `Blasi`. This implies that even for these problems there is a substantial fraction of the runs which, although a global optimisation was performed, did not provide the global optimum. Assessment of optimality gaps for individual optimisation methods revealed that `MS+CMA-ES`, followed by `MS+fides` and `SaCeSS+Ipopt` achieved the lowest medians, while `SaCeSS+Ipopt` provided the best worst-case results (Figure 4E). Complementarily, when comparing the numbers of problems solved by the best and the worst runs of each optimisation method, we found that the two best performing ones in terms of best run solved only two problems fewer in their worst run, indicating good reliability (Supplementary Figure S5). For other optimisation methods, the difference in the number of solved problems in the best and worst runs was as high as 10. Indeed, there was good agreement between the median optimality gap achieved by algorithms and the number of benchmark problems it was able to solve.

To study the impact of the choice of globalisation and parallelisation strategy, we compared `SaCeSS+Ipopt` with `SaCeSS` alone and with multi-start optimisation using `Ipopt` (namely `MS+cyipopt`). `SaCeSS+Ipopt` achieved more reliable results than either component alone: the median number of problems solved across the 10 runs increased from 18 for `MS+cyipopt` and 19 for `SaCeSS` to 23 for `SaCeSS+Ipopt`, suggesting a synergistic effect of combining `SaCeSS`-based global search with local refinement (Supplementary Figure S5).

In summary, we found that some optimisation methods clearly outperformed others in finding the optimal solution on the parameter estimation benchmarks considered. `MS+fides` and `MS+CMA-ES` solved the largest number of problems. `MS+fides`, `MS+CMA-ES`, and `SaCeSS+DHC` made up the smallest set sufficient to solve all 30 problems. The different `SaCeSS` variants ranked high in the number of solved problems. Among these, `SaCeSS+Ipopt` provided the best median optimality gap and reliability.

### Minimal resource requirement for optimisation

The computational resources required to solve optimisation problems can be substantial. Accordingly, an assessment of the minimal required resources is critical. Here, we assessed the resource requirements by determining the minimal time an optimisation method required for reaching an acceptable objective function value according to a *χ*^2^-based rejection test with a significance level of 0.05, corresponding to minimal optimality gap of 1.92 compared to the observed global best value (Figure 5A). We denote this time as *first hitting time* and evaluate it for each combination of benchmark problem and optimisation method. As all optimisation methods used the same computational infrastructure, the first hitting time provides information about the relative resources required to achieve a good fit.

**Fig 5:**
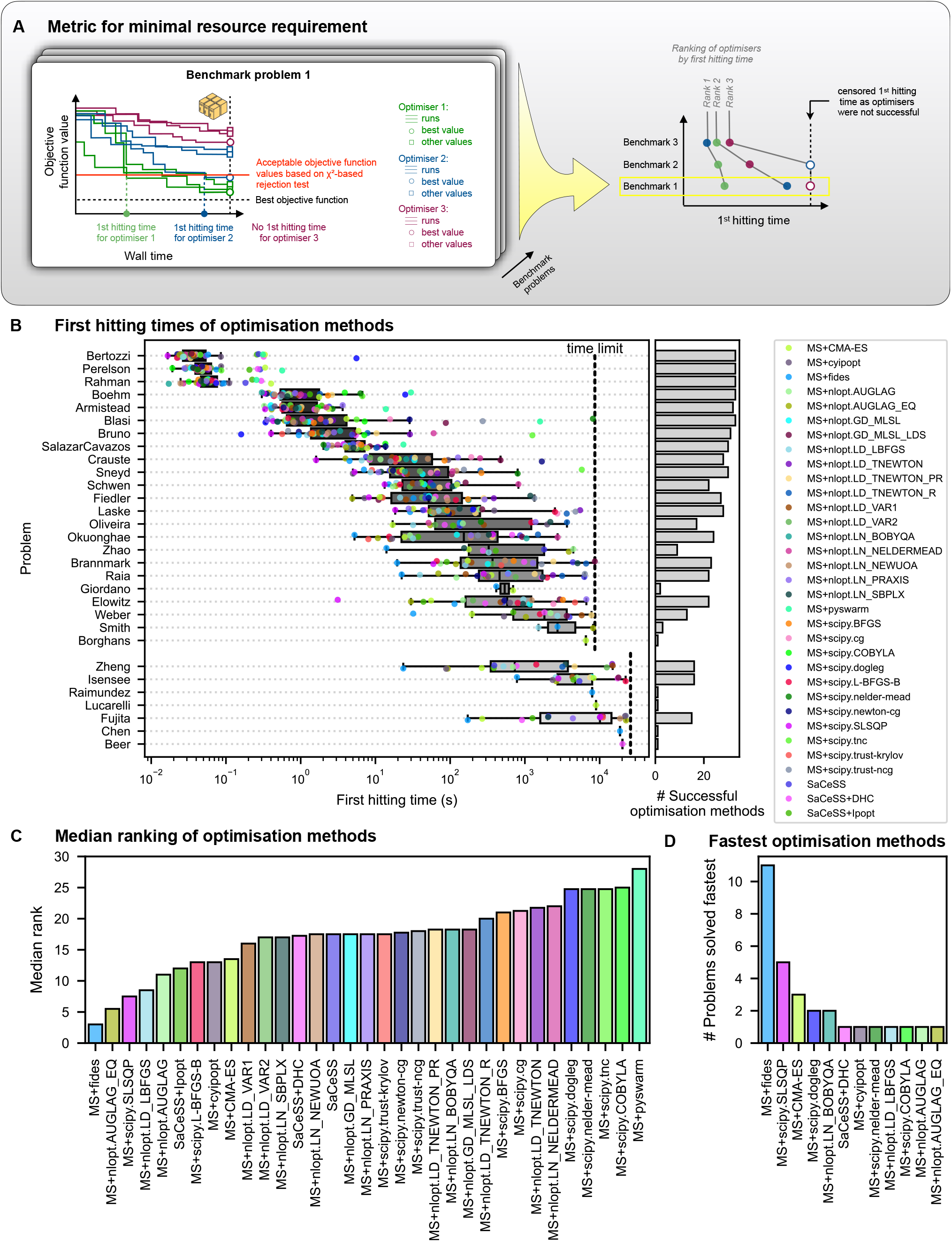
Resource requirement analysis. (A) Illustration of the objective function values for different optimisation runs as a function of wall time, and an illustration of the first hitting time. Here, the first hitting time is the earliest time point the optimality gap falls below 1.92, which corresponds to a *χ*^2^-based rejection test with significance level 0.05. (B) First hitting times for all problem-method-pairs (minimum across 10 runs) and the number of successful optimisation methods per problem. Problems are sorted by the median first hitting time. (C) Median rank of optimisation methods in first hitting time analysis. In case of ties, the average rank was used. (D) Number of times an optimisation method was the fastest in the first hitting time analysis. The reference points for the optimality gaps were the best values achieved by any run of the optimisation methods shown in (C) on Marvin.

Our analysis of the first hitting time revealed that the considered benchmark problems require vastly different computational resources (Figure 5B). While the benchmark problems by Bertozzi et al. (2020) and Perelson et al. (1996) could be solved by most optimisation methods in below one second, several challenging problems required at least one hour. Unsurprisingly, the number of successful optimisation methods decreased as the median first hitting time increased. This indicates that, in the end, whether an optimisation method converges depends on how efficiently it exploits the available computational resources.

From the first hitting times we constructed a ranking, with the fastest optimisation method indicated as rank 1 and the method that did not solve the problem with rank 33 (in case of ties, the average rank was used). The assessment of the ranking revealed that `MS+fides` achieved lowest median rank, followed by `MS+nlopt.AUGLAG_EQ`, `MS+scipy.SLSQP`, `MS+nlopt.LD_LBFGS`, `MS+nlopt.AUGLAG`, and `SaCeSS+Ipopt` (Figure 5C). Hence, multi-start optimisation with advanced, gradient-based local optimisation algorithms can surpass enhanced scatter search optimisation methods. Indeed, `MS+fides`, which was able to solve the highest number of problems, was for 11 benchmark problems the method with the minimal first hitting time. As the best competitors provide minimal first hitting time for at most five benchmark problems, `MS+fides` is a good candidate if time is restrictive (Figure 5D).

Interestingly, `MS+CMA-ES`, the method which solved the second-most of the optimisation problems, did not achieve very low first hitting times, implying that it requires more resources. However, `MS+CMA-ES` was still the highest-ranking derivative-free method in this analysis.

In summary, we found that optimisation problems in our benchmark require vastly different computational resources, with first hitting times ranging from below one second to more than one hour. Multi-start local optimisation with advanced gradient-based algorithms, in particular `MS+fides`, achieved the lowest median first hitting time and was the fastest method on most benchmark problems.

### Efficiency of optimisation

A successful optimisation run is an important first step; yet good scientific practice requires reproducibility. Hence, we studied how efficiently the methods generated independent successful optimisation starts from random initial points. For each optimisation method *i* and benchmark problem *j*, we counted the number of successful starts, 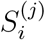, across the 10 runs, using the same optimality gap threshold as above (Figure 6A). If all methods are evaluated under the same problem-specific compute budget, this count directly reflects how efficiently a method converts the available resources into successful independent starts. While single-start methods, such as `SaCeSS`, can generate at most one successful start per run, multi-start methods can produce multiple successful starts within the same budget.

**Fig 6:**
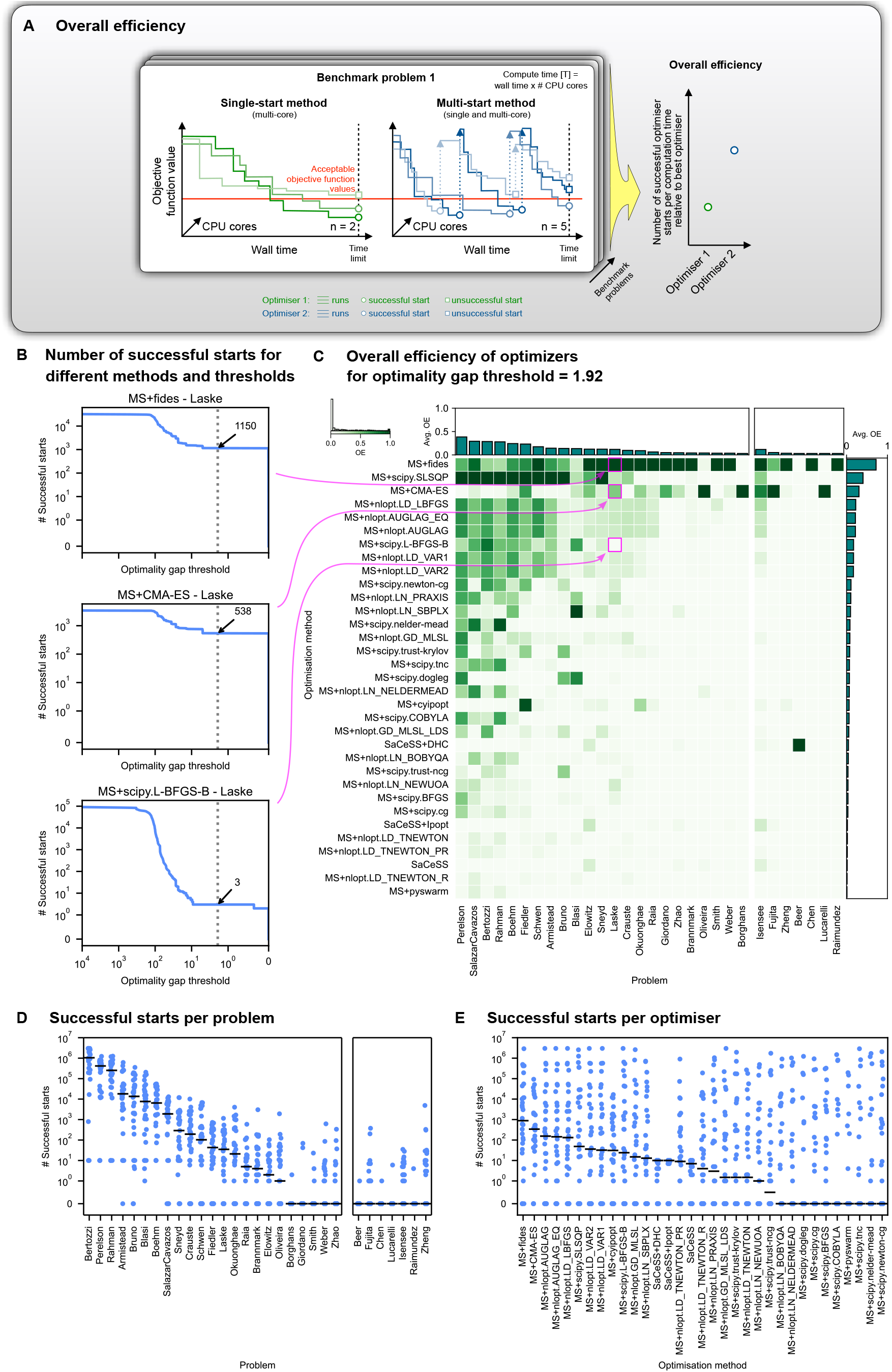
Overall efficiency analysis. (A) Illustration of the metric for overall efficiency for single- and multi-start methods. It is shown how the number of successful starts is, for a given threshold, determined based on the optimisation trajectories. (B) For three exemplary combinations of problem and optimisation method, the number of successful starts at different optimality gap thresholds (sum across the 10 optimisation runs). The dashed lines indicate the threshold applied to generate subfigure (C), the numbers next to the arrow indicate the number of successful starts at that threshold. (C) Heatmap of overall efficiencies for combinations of benchmark problems and optimisation methods, as well as bar plots for the average overall efficiency per benchmark problem and method. (D) Number of successful starts for the different benchmark problems. Each point represents the sum of successful starts across the 10 runs of one optimisation method. The black lines indicate the medians. (E) Number of successful starts for the different optimisation methods. Each point represents the sum of successful starts across the 10 runs for one problem. The black lines indicate the medians. The reference points for the optimality gaps were the best values achieved by any run of the optimisation methods shown in (E) on Marvin.

To account for differences in problem complexity, we normalised the number of successful starts for each benchmark problem by the best-performing method on that problem and defined the overall efficiency for runs with equal time as

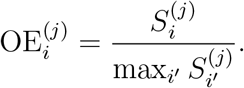

To account for possible differences in compute time between runs, 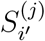 was itself normalised by compute time (see Materials and Methods). By construction, the optimisation method producing the largest number of successful starts for a given problem has an overall efficiency of one, whereas methods producing fewer successful starts achieve lower values. Methods without successful starts have an overall efficiency of zero. As this measure is scale-free, it can be averaged across benchmark problems.

Our assessment reveals that gradient-based multi-start optimisation methods achieved, for most benchmark problems the best overall efficiency (Figure 6B&C). Indeed, four out of the top five methods used gradient-based local optimisation algorithms, which exploit first- or first- and second-order information. The optimisation method with the highest overall efficiency was `MS+fides`, followed by `MS+scipy.SLSQP`, `MS+CMA-ES` (the only gradient-free method among the top five), `MS+nlopt.LD_LBFGS`, and `MS+nlopt.AUGLAG_EQ`. Interestingly, `MS+fides` and `MS+scipy.SLSQP` seemed to favour distinct subsets of problems. Whereas `MS+scipy.SLSQP` achieved highest overall efficiencies for problems with below 10 estimated parameters, `MS+fides` performed better on higher-dimensional problems (Figure 6C and Supplementary Table S2).

Single-start methods, namely `SaCeSS` with and without a local optimiser, achieved mostly low overall efficiencies in our benchmarking. High overall efficiencies were only observed for benchmark problems that were not reproducibly solved by multi-start optimisation methods.This is to be expected, as the considered single-start methods do not employ sophisticated stopping criteria but simply exploit the available resources. In particular for simpler problems, a reduction in the wall-time limit would improve the overall efficiency of single-start methods, while having little influence on the overall efficiency of multi-start methods. In the current setup, we found that `MS+fides` achieved, at the median, almost a hundred times more successful starts than single-start methods (Figure 6E).

The assessment on a per-model basis indicated that both the overall efficiency (Figure 6C) and the number of successful starts (Figure 6D) vary over several orders of magnitude. For a few benchmark problems, more than 10^6^ successful starts were observed, whereas for others we observed, across optimisation methods, only a single successful start. For the latter, the reproducibility and reliability of the best fit may still be an issue.

In summary, the overall efficiency analysis shows that multi-start strategies are advantageous when reproducibility is to be assessed through independent successful optimiser starts. Under a fixed compute budget, they can generate multiple independent solutions, whereas single-start strategies can produce at most one successful start per run. Single-start methods should therefore be combined with effective stopping or restart criteria to avoid inefficient use of computational resources, particularly for simpler problems.

### Novel optimisation method combinations

The benchmark results might provide the basis for a data-driven development of optimisation methods. To assess this, we followed up on our earlier hypothesis of a potential synergy between hybrid global derivative-free and local gradient-based methods as observed for `SaCeSS+Ipopt` (Supplementary Figure S5). The identification of `fides` as the best local algorithm suggested an assessment of a `SaCeSS` variant that uses `fides` as local solver, yielding `SaCeSS+fides`. As the original Fortran/`C` implementation of `SaCeSS` could not be easily extended to incorporate alternative local optimisation algorithms, we developed a single-node, multiprocessing Python implementation which we refer to as `pySaCeSS` in this manuscript (Figure 7A). `pySaCeSS` features a modular design that supports any optimisation algorithm interfaced by `pyPESTO` as a local search method.

**Fig 7:**
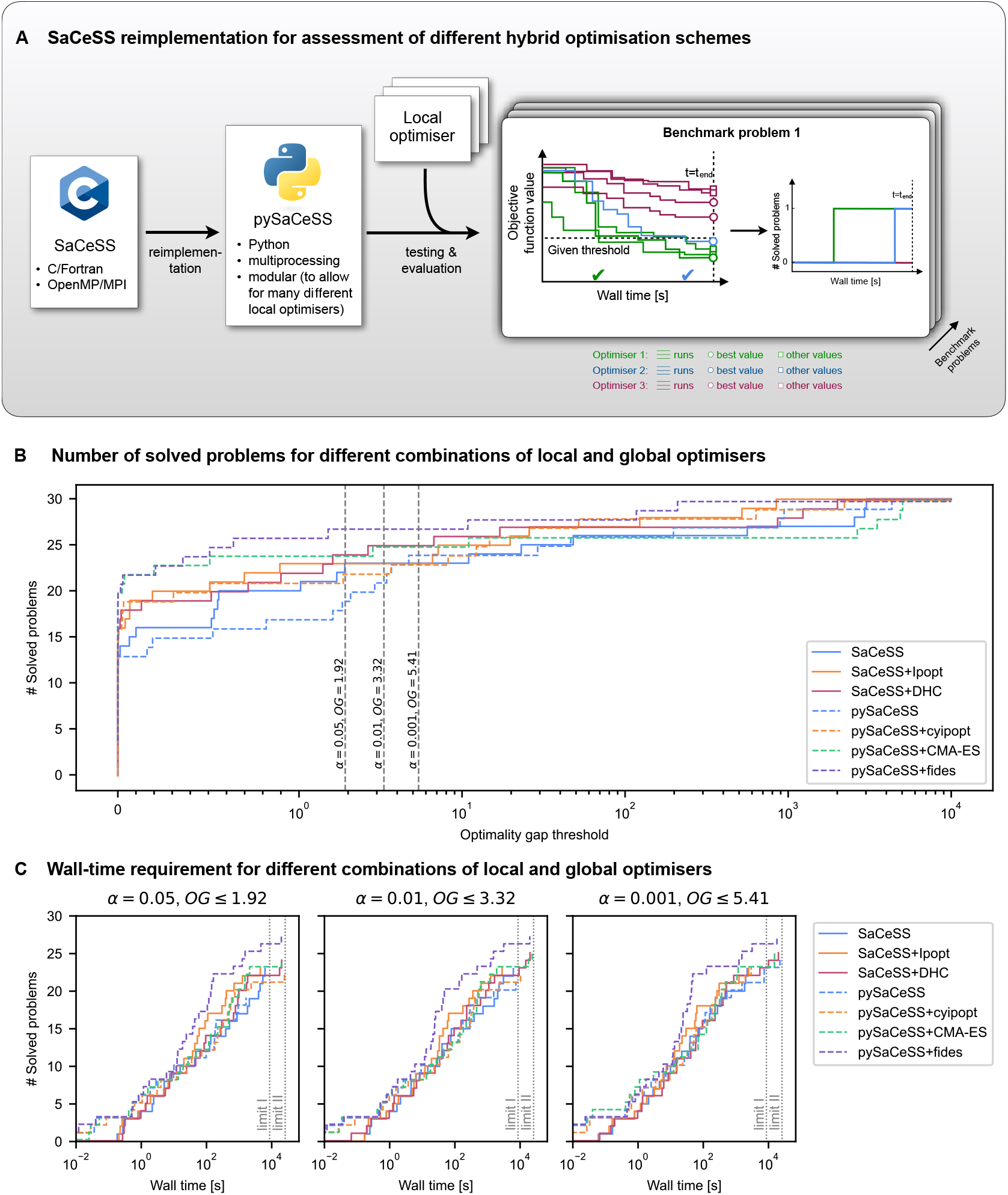
Analysis of novel optimisation model developed based on benchmarking. Overview of the two SaCeSS implementations. (B) Number of solved problems for different optimisation methods across optimality gap thresholds. Thresholds for three significance levels are highlighted. The best final objective value across 10 runs is used. (C) Performance profiles illustrating the number of solved problems over time for each method under three optimality gap thresholds corresponding to significance levels of 0.05, 0.01 and 0.001 in a *χ*^2^-based rejection test. The best value across the 10 runs is used at each timepoint. The lines are offset slightly to reduce overplotting. The reference points for the optimality gaps were the best values achieved by any run of the optimisation methods shown in Figures 1 and 7 on Marvin.

A comparison of `SaCeSS` and `pySaCeSS`, both, without a local solver and with `Ipopt` as local solver, showed reasonable agreement. Without a local solver, `SaCeSS` solved 23 benchmark problems across all three considered significance levels (*α* = 0.05, 0.01 and 0.001), whereas `pySaCeSS` solved 19, 21 and 24 problems, respectively (Figure 7B). This indicated that the original Fortran/C `SaCeSS` version (Penas et al., 2017) achieves slightly better performance than the reimplementation. Indeed, at the *α* = 0.05 significance level, it solved four more problems than the reimplementation (Supplementary Figure S7A&B); in combination with `cyipopt`/`Ipopt`, the gap was only one problem.

We then used the modular reimplementation to benchmark enhanced scatter search in combination with several top-performing local optimisation algorithms. As suggested by our benchmarking results, `pySaCeSS+fides` outperformed or was on par with all other combinations of `SaCeSS` and `pySaCeSS`, with and without local solvers. For most optimality gap thresholds, it is indeed the top performing method in terms of solved problems (Figure 7B). The assessment of convergence speed revealed that `pySaCeSS+fides` consistently outperformed the other considered configurations (Figure 7C). While `pySaCeSS+CMA-ES` was on par with `pySaCeSS+fides` at very small optimality gap thresholds (*<* 0.5), it solved fewer problems at higher thresholds and showed slower convergence.

In summary, the modular reimplementation of `SaCeSS`, `pySaCeSS`, improves access to and flexibility of the underlying algorithm. This enables systematic testing of additional local optimisation algorithms and hyperparameters. Given the large number of possible choices, the benchmarking results presented here can be used to prioritise future evaluations.

## Discussion

Maximum likelihood and maximum a posteriori parameter estimation for dynamical models requires reliable and efficient optimisation methods. Over the past decades, a plethora of approaches has been developed, yet clear recommendations are lacking and some methods are difficult to apply in practice. To address this gap, we implemented a comprehensive, standardised pipeline and used it to benchmark published parameter estimation problems for ODE models in systems biology, medicine and epidemiology.

For the benchmark problems from the application domains, our results show that some optimisation methods consistently outperform others. Our study yields several clear, practically relevant conclusions:

1. To maximise the probability of finding the best objective function value, the multi-start methods `MS+fides` and `MS+CMA-ES` were the most reliable methods for genuinely challenging problems. Close competitors were `MS+nlopt.AUGLAG_EQ` (using `MS+nlopt.LD_LBFGS` internally), and the single-start methods `SaCeSS+DHC`, `SaCeSS`, and `SaCeSS+Ipopt`.
2. To obtain parameter estimates in a time-critical setting, `MS+fides` is the method of choice. It is comparatively reliable and achieves favourable first-hitting times, most likely due to its efficient use of gradient information.
3. To ensure reproducibility via multiple independent optimisation starts, the choice depends on the problem complexity. If a problem can be solved with multi-start methods, `MS+fides` is preferable owing to its high efficiency and advanced stopping criteria. However, we encountered problems that were solved only by `SaCeSS+DHC` or `MS+CMA-ES`, possibly because of the high cost of gradient evaluations relative to objective function evaluations for these problems.

To arrive at these conclusions, we considered a massive amount of real world benchmark problem optimisation method pairs. This is a completely new scale, far exceeding previously available small scale studies with limited problem and optimiser spectrum. The reliability was ensured by performing the analysis on two different HPC systems at two different locations, using in total over 1.5 million core-hours.

Clear recommendations and guidelines are valuable, yet their practical relevance depends on easy availability of the recommended methods. While several algorithms were already available in user-friendly packages, we here provide a Python reimplementation of `SaCeSS`, which can be seamlessly used within the widely adopted `AMICI`/`pyPESTO` workflow (Schälte et al., 2023).

The modular nature of the reimplementation also enabled us to assess different method combinations. This revealed that combining an advanced global search strategy with the best-performing local solver, i.e. `pySaCeSS+fides`, can further improve performance. We regard this not as a final outcome, but as the starting point for further work. Among other directions, it would be worthwhile to equip `SaCeSS` with alternative stopping criteria to improve computational efficiency without sacrificing robustness.

In addition to comparing optimisation methods, we also investigated the relative difficulty of the benchmark problems themselves. Some problems turned out to be comparatively easy, while others were considerably harder. We explored the possibility of training a statistical model to predict optimiser-specific performance characteristics from simple model properties, such as the number of parameters or data points. Unfortunately, the predictive power of this model was limited, suggesting that either the dependence on such simple characteristics is weak, or that the current set of benchmark problems is too small to extrapolate reliably to new models. Expanding the collection of PEtab benchmark problems would therefore be valuable; to facilitate this, we are in the process of simplifying adoption of PEtab (Jost et al., 2026).

This study has several limitations: First, we did not include all algorithms available in `NLopt` and `scipy`, mostly due to constraints such as upper bounds on the number of optimisation variables. Moreover, a small number of solvers do not support both lower and upper bounds. While this can in principle be addressed by parameter transformations, here we opted for a subsequent filtering step, which may yield suboptimal outcomes for some methods. Second, we considered forward and adjoint sensitivities, but decided before the optimisation on the method based on computation time and success rate. In theory, it is possible that a sensitivity computation method with disadvantageous sensitivity computation time or success rate might still yield a better optimisation performance, but we consider this unlikely. We previously found a good agreement of sensitivity computation and optimiser performance (Villaverde et al., 2018). Third, our results depend on the chosen computation time budget. One could, in principle, use the same maximal wall time for all problems to simplify interpretation.

However, for short budgets even more of the more complex problems would remain unsolved or strongly affected by stochasticity, whereas very long budgets would waste computational resources on simpler problems. We therefore opted for problem-specific time limits that balance these considerations.

Furthermore, we note that the results necessarily depend on the hyperparameters, such as termination criteria, used for the individual optimisation algorithms. In this study, we used default hyperparameters whenever possible to obtain a realistic and reproducible assessment of out-of-the-box performance. However, these defaults are not necessarily optimal for the benchmark problems considered here, and they can influence the outcome of all metrics. For example, several local optimisation algorithms use stopping tolerances in the order of 10^−6^. Such thresholds are meaningful in many applications, but they can prevent further improvement once the objective function value is already close to the best value found. Consequently, some methods may achieve worse ranks in analyses based on very small differences in the final objective value, not because they fail to identify the relevant optimum, but because they terminate earlier than methods with stricter stopping criteria. Conversely, global and hybrid methods often use the full available wall time unless equipped with dedicated stopping rules. This affects not only final solution quality but also efficiency measures. For instance, `SaCeSS` uses the allocated resources cooperatively for a single start, whereas multi-start methods split the same resources into many independent starts. For difficult problems this can improve robustness, but for simpler problems the same total resources would likely be used more efficiently by splitting the computation into several independent `SaCeSS` runs or by introducing more aggressive stopping criteria, yielding improved overall efficiency.

Additionally, optimisation performance can heavily depend on ODE solver hyperparameters, such as integration tolerances, sensitivity analysis method, or the ODE solver as such. The exact choice of hyperparameters can have a strong impact on the relative cost of a gradient evaluation compared to an objective evaluation, thereby biasing the results toward gradient-based or gradient-free optimisation methods, or they can heavily influence the failure rate, thereby, biasing the results towards methods with better error handling. Here, we used the default settings of the `AMICI` simulator, which may not be optimal settings for a given combination of optimisation method and problem and may render certain optimisations more challenging than they could be. Accordingly, the ranking of optimisation problems may very much depend on the applied ODE solvers or options.

In summary, our benchmarking emphasises that the choice of optimisation method can have a substantial impact on reliability, efficiency and reproducibility in parameter estimation for dynamical biological models. By identifying well-performing optimisation methods, and by providing an accessible, modular implementation of `SaCeSS` that integrates with established workflows, we offer concrete, data-driven guidance for practitioners. We anticipate that these results, together with an expanded set of benchmark problems, will support more robust parameter estimation and help to standardise optimisation practices in systems biology, medicine and epidemiology.

## Materials and Methods

### Mathematical models and experimental data

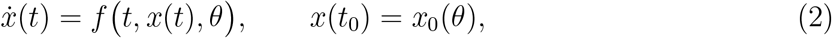

with parameter vector 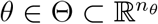. Initial conditions may depend on *θ* and, when not known *a priori*, and not implicitly specified as steady-state constraint, are estimated alongside other parameters. Experimental conditions might alter the initial state or the vector field, but for readability, we present here only the case for a single experimental condition.

Experimental readouts are described by observables

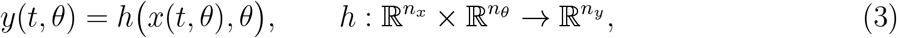

evaluated at measurement time points *t*_*k*_, *k* = 1, …, *n*_*t*_. We consider different noise models, including the Gaussian measurement noise,

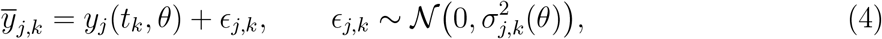

for observables *j* = 1, …, *n*_*y*_. The measured data are collected in a dataset 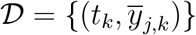, potentially across multiple experimental conditions and replicates, specified via the PEtab format (Schmiester et al., 2021).

### Optimisation problems

Maximum likelihood or maximum a posteriori estimation problems were considered. Under the Gaussian noise assumption, the likelihood of the data *D* given parameters *θ* is

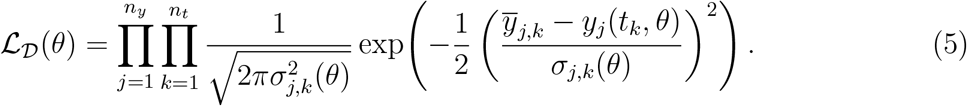

We minimise the negative log-likelihood

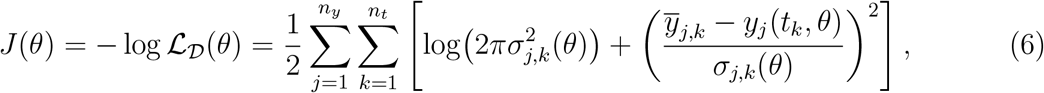

or the (non-normalized) negative log-posterior (*J*(*θ*) = −(log *L*_*D*_(*θ*) +log *p*(*θ*)), with *p*(*θ*) denoting the prior distribution), subject to the ODE dynamics (2) and simple bound constraints *θ*^*L*^ ≤ *θ* ≤ *θ*^*U*^. The optimal parameter vector

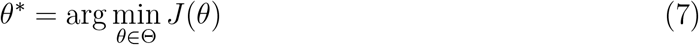

was found via numerical optimisation. We used a single-shooting approach in which, for each candidate *θ*, the ODE system is integrated numerically and the objective *J*(*θ*) is evaluated.

### Metrics for benchmarking

#### Overall efficiency (OE)

To compare efficiency across methods and problems we used the overall efficiency measure introduced in Villaverde et al. (2018). For optimisation method *i* on benchmark problem *j*, we determined the overall computation time 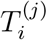 and the number of successful starts 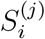 across all 10 runs, yielding the average computation time that the *i*-th optimisation method required to produce a successful start as

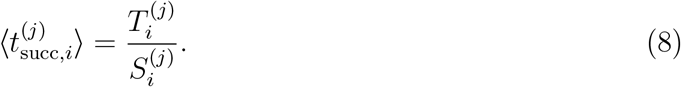

The overall efficiency on problem *j* is then

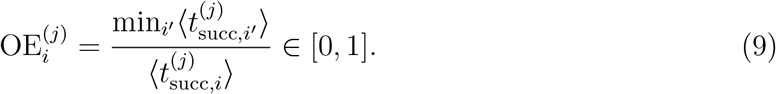

The best optimisation method for a given problem has 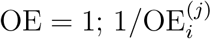 indicates how many times longer method *i* must be run, relative to the best method, to obtain one successful start.

### Implementation

#### Benchmark problems

We benchmarked optimisation methods using 30 parameter estimation problems from the PEtab benchmark collection (Schmiester et al., 2021) (https://github.com/Benchmarking-Initiative/Benchmark-Models-PEtab, Release 2024-11-11). The collection overlaps with the problems studied in Hass et al. (2019), but includes additional problems, e.g., for the spread of infectious diseases. The collection covers a broad range of characteristics, including numbers of parameters, state variables, experimental conditions and data points, as well as different input and noise models. Models in the benchmark collection which contained reported issues or which required more than 10 seconds for a single objective function evaluation were excluded. Based on prior experience and exploratory runs, we classified the problems into two subsets: subset I (comparatively easy) and subset II (more challenging with respect to required optimisation time) (Supplementary Table S2). This classification was used solely to allocate computational resources.

#### Optimisation frameworks and algorithms

We benchmarked 33 different optimisation methods (Figure 1A). The employed package versions are listed in Supplementary Table S1. The different optimisation algorithms were interfaced through three optimisation frameworks tailored to PEtab problems:

- `pyPESTO` (Schälte et al., 2023): a Python-based parameter estimation toolbox implementing multi-start local optimisation parallelised over starting points. It interfaces with AMICI (Fröhlich et al., 2021) for simulation and sensitivities and provides access to a wide range of local and global solvers, including `nlopt` (Johnson, 2023), `scipy` (`scipy.optimize.minimize`) (Virtanen et al., 2020), `CMA-ES` (Hansen et al., 2019), `fides` (Fröhlich and Sorger, 2022) and `Ipopt` (Wächter and Biegler, 2006). We note that some solvers from `nlopt` and `scipy` were excluded, for example, due to applicability constraints and high failure rates in initial tests.
- `parPE` (Schmiester et al., 2020): a `C++` library for parameter estimation of PEtab problems. `parPE` interfaces, among others, `Ipopt` and uses AMICI for model simulation and sensitivity analysis.
- `SaCeSS` (Penas et al., 2017): the Self-adaptive Cooperative enhanced Scatter Search, an island-based evolutionary global optimisation framework built around enhanced Scatter Search (Rodríguez-Fernández et al., 2006; Egea et al., 2010; Villaverde et al., 2012). Independent eSS instances (“islands”) run in parallel, and a manager process manages asynchronous communication and hyperparameter adaptation. We extended `SaCeSS` to support PEtab problems by interfacing `parPE`.

Some of the selected optimisation methods did not support box constraints. Therefore, optimisation results were filtered for admissible/feasible parameter values (satisfying the box constraints) during post-processing. Function evaluations with inadmissible parameters are still considered in the compute times.

All optimisation methods were run with their default hyperparameters, apart from the wall time limit. For the `nlopt` methods `nlopt.{GD_MLSL,GD_MLSL_LDS,AUGLAG,AUGLAG_EQ}`, which require specifying an inner optimisation algorithm, `nlopt.LD_LBFGS` was used as per `pyPESTO`’s default settings.

For multi-start methods, the number of starts was determined implicitly by how many fits could be executed within the allocated wall time, unless the number of starts exceeded 300,000, in which case the respective run was terminated immediately. `SaCeSS`-based methods used the wall time limit as the sole exit criterion; they were run with a number of workers equal to the number of allocated cores minus one. For each run, we recorded the best objective value achieved as a function of wall-clock time, enabling fair comparisons of convergence behaviour and efficiency under identical resource constraints.

#### Model simulation

In all cases, `AMICI` (Fröhlich et al., 2021) was used for model simulation, and where required by derivative-based optimisation methods, for sensitivity analysis. Gradients were computed using forward or adjoint sensitivities (Supplementary Table S2). For optimisation algorithms that strictly required the Hessian of the objective function (`scipy.{dogleg,trust-ncg,trust-krylov}`), the Hessian was approximated via the empirical Fisher information matrix constructed from first-order forward sensitivities. For simulation hyperparameters, such as integration tolerances, we used `AMICI`’s default settings.

#### High-performance computing setup

All computations were performed on two high-performance computing (HPC) infrastructures:

- **FinisTerrae III** (https://www.cesga.es): CPU nodes with 2× Intel Xeon “Ice Lake” 8352Y processors (32 cores at 2.2 GHz, 64 cores per node) and 256GB RAM, connected via Infiniband HDR100.
- **Marvin** (https://www.hpc.uni-bonn.de): CPU nodes with 2× Intel Xeon “Sapphire Rapids” Platinum 8468 processors (48-core/96-thread 2.10 GHz, 96 cores per node) and 1024GB DDR5 4800MHz RAM, connected via Mellanox Infiniband NDR 200 Gbit/s.

The computations were containerized to maximize reproducibility and to facilitate reuse. We used identical MPI implementations and compiler settings on both systems and replicated key experiments across the two clusters to verify robustness. We confirmed that containerization with `apptainer` only had a negligible impact on the run time (*<* 5%).

For subset I problems we allocated 12 cores for 3 h per optimiser–problem combination; for subset II we allocated 24 cores for 9 h (Supplementary Table S2). Each optimisation method was run 10 times per problem and supercomputer to account for stochastic variability. Processes were killed at the end of the allocated time budget and the progress of the still-running multi-start starts were discarded. After executing all runs, we noticed that there was a non-negligible number of runs for which the last start that was killed accounted for up to 20% of the total budget. To conserve computational resources without unfairly penalising long-running optimisation algorithms, we performed the analysis on a retrospectively reduced wall time budget of 80% of the original allocation.

#### SaCeSS reimplementation in Python

To facilitate the evaluation of `SaCeSS` in combination with additional local optimisation algorithms, we reimplemented a basic `SaCeSS` version in Python (only shared-memory parallelism, only continuous variables). The Python implementation enables combining the `SaCeSS` method with any of the optimisation algorithms interfaced by `pyPESTO`, e.g., those from the `scipy` or `nlopt` packages. The code is available at https://github.com/ICB-DCM/pyscat/ (class `SacessOptimizer`). We refer to this implementation as `pySaCeSS` in this study.

## Data availability statement

Data and data analysis code, as well as containers used for data generation, are available online at https://github.com/ICB-DCM/optimizer-benchmark-2026-suppl-code-and-data/ and https://doi.org/10.5281/zenodo.21296723.

## Acknowledgments

The authors gratefully acknowledge (1) the University of Bonn for providing access to the Marvin HPC cluster, and (2) the CESGA (Centro de Supercomputación de Galicia) for providing access to its FinisTerrae III supercomputer.

## Author contributions

D.R.P. and S.G. conducted the benchmark study on the two HPC infrastructures located at CESGA and at the University of Bonn, respectively. D.W. implemented the Python version of SaCeSS and the PEtab interface for parPE, analysed the data, and supported the execution of the benchmark study. P.L. assisted with the visualisation of the fitting results. J.R.B. and J.H. conceived, conceptualised and guided the study process. S.G., J.R.B., D.R.P., and J.H. authored the first version of the manuscript. All authors reviewed and revised the manuscript.

