## Supplementary Figures and Tables for "Large-scale analysis of optimisation methods for parameter estimation problems in the life sciences"

**Supplementary Table S1.** Employed software packages used for simulation and optimisations in this study, with version identifier, and reference to implementation or appearance in literature. For pyPESTO, some adaptations related to the result storage format were necessary—details are provided in the supplementary data.

| Software | Version | Link | Reference |
|---|---|---|---|
| **AMICI** | 0.30.1 | `https://github.com/AMICI-dev/AMICI` | [1] |
| **apptainer** | 1.4.2-1.el9 | `https://apptainer.org` | [2] |
| **CMA-ES** | 4.0.0 | `https://github.com/CMA-ES/pycma` | [3] |
| **cyipopt** | 1.5.0 | `https://github.com/mechmotum/cyipopt` | [4] |
| **fides** | 0.7.8 | `https://github.com/fides-dev/fides/` | [5] |
| **Ipopt** | 3.14 | `https://github.com/coin-or/Ipopt` | [6] |
| **nlopt-python** | 2.8.0 | `https://github.com/DanielBok/nlopt-python/` | [7, 8] |
| **parPE** | 0.7.0 | `https://github.com/ICB-DCM/parPE` | [9] |
| **PEtab** | 0.5.0 | `https://github.com/PEtab-dev/libpetab-python` | [10] |
| **pyPESTO** | 0.5.4 (modified) | `https://github.com/ICB-DCM/pyPESTO` | [11] |
| **pyscat** | 0.0.1.post2.dev24+g03f949dee | `https://github.com/ICB-DCM/pyscat/` | [12] |
| **pyswarm** | 0.6.0 | `https://github.com/tisimst/pyswarm` | [13] |
| **SaCeSS** | 1.0.2 | `https://github.com/davidrpenas/sacess_petab` | [14] |
| **SciPy** | 1.15.2 | `https://github.com/scipy/scipy` | [15] |

**Supplementary Table S2. Main properties of the considered benchmark problems.** Various model dimensions, employed sensitivity method, and original reference for each problem. Problems were assigned to two subsets, I and II, with different CPU budgets per run (see main text for details). In case of derivative-free optimisers, no sensitivities were computed; for optimisers accepting gradients, the indicated method was applied (ASA: adjoint sensitivity analysis; FSA: forward sensitivity analysis); for optimisers requiring the Hessian, FSA was used for all problems. Estimated parameters, observables, and simulation conditions are counted as defined in the PEtab implementation of each problem. State variables are counted from the `AMICI`-imported model.

| ID | # Estimated parameters | # State variables | # Observables | # Conditions | Subset | Sensitivity method | Citation |
|---|---|---|---|---|---|---|---|
| **Armistead** | 14 | 4 | 4 | 2 | I | FSA | [16] |
| **Beer** | 72 | 4 | 2 | 19 | II | ASA | [17] |
| **Bertozzi** | 8 | 3 | 2 | 2 | I | FSA | [18] |
| **Blasi** | 9 | 16 | 15 | 1 | I | ASA | [19] |
| **Boehm** | 9 | 8 | 3 | 1 | I | FSA | [20] |
| **Borghans** | 23 | 3 | 1 | 1 | I | FSA | [21] |
| **Brannmark** | 22 | 9 | 3 | 8 | I | ASA | [22] |
| **Bruno** | 13 | 7 | 5 | 6 | I | FSA | [23] |
| **Chen** | 155 | 500 | 3 | 4 | II | ASA | [24] |
| **Crauste** | 12 | 5 | 4 | 1 | I | FSA | [25] |
| **Elowitz** | 21 | 8 | 1 | 1 | I | FSA | [26] |
| **Fiedler** | 22 | 6 | 2 | 3 | I | FSA | [27] |
| **Fujita** | 19 | 9 | 3 | 6 | II | FSA | [28] |
| **Giordano** | 50 | 10 | 7 | 1 | I | ASA | [29] |
| **Isensee** | 46 | 25 | 3 | 123 | II | ASA | [30] |
| **Laske** | 13 | 34 | 13 | 3 | I | FSA | [31] |
| **Lucarelli** | 84 | 33 | 65 | 16 | II | ASA | [32] |
| **Okuonghae** | 16 | 8 | 2 | 1 | I | FSA | [33] |
| **Oliveira** | 12 | 9 | 2 | 1 | I | ASA | [34] |
| **Perelson** | 3 | 4 | 1 | 1 | I | FSA | [35] |
| **Rahman** | 9 | 7 | 1 | 1 | I | FSA | [36] |
| **Raia** | 39 | 14 | 8 | 4 | I | FSA | [37] |
| **Raimundez** | 136 | 22 | 79 | 170 | II | ASA | [38] |
| **SalazarCavazos** | 6 | 75 | 3 | 4 | I | ASA | [39] |
| **Schwen** | 30 | 11 | 4 | 19 | I | FSA | [40] |
| **Smith** | 25 | 109 | 9 | 35 | I | ASA | [41] |
| **Sneyd** | 15 | 6 | 1 | 9 | I | FSA | [42] |
| **Weber** | 36 | 7 | 8 | 2 | I | ASA | [43] |
| **Zhao** | 28 | 4 | 1 | 7 | I | FSA | [44] |
| **Zheng** | 46 | 15 | 15 | 1 | II | ASA | [45] |

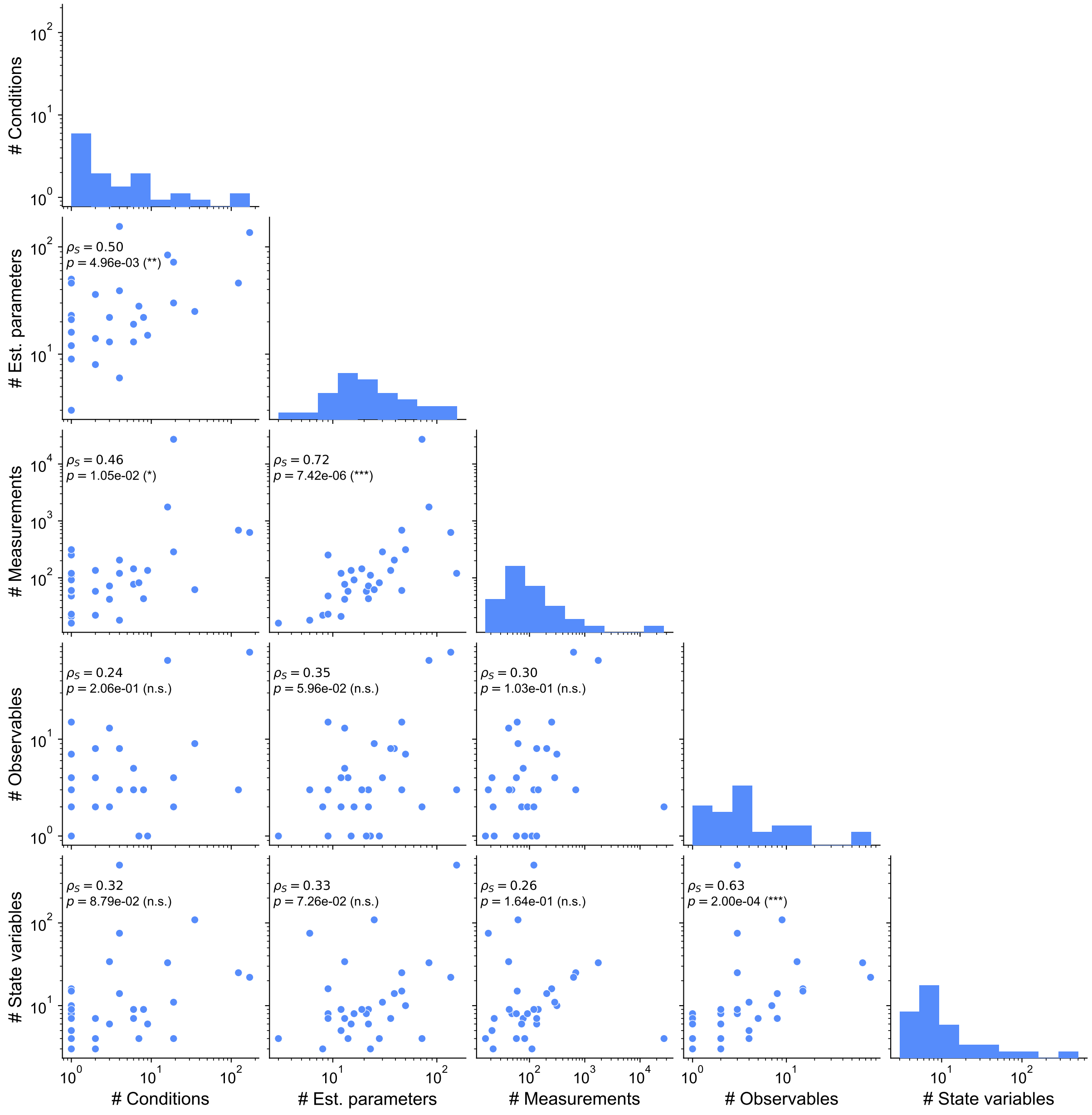

**Supplementary Figure S1. Characteristics of the selected benchmark problems.** The number of state variables, observables, estimated parameters, measurements and experimental conditions is visualised: (Diagonal) Histograms for individual characteristics; (Off-diagonal) Scatter plots for pairs of model characteristics. The Spearman correlations $\rho_S$ and their $p$-values are indicated.

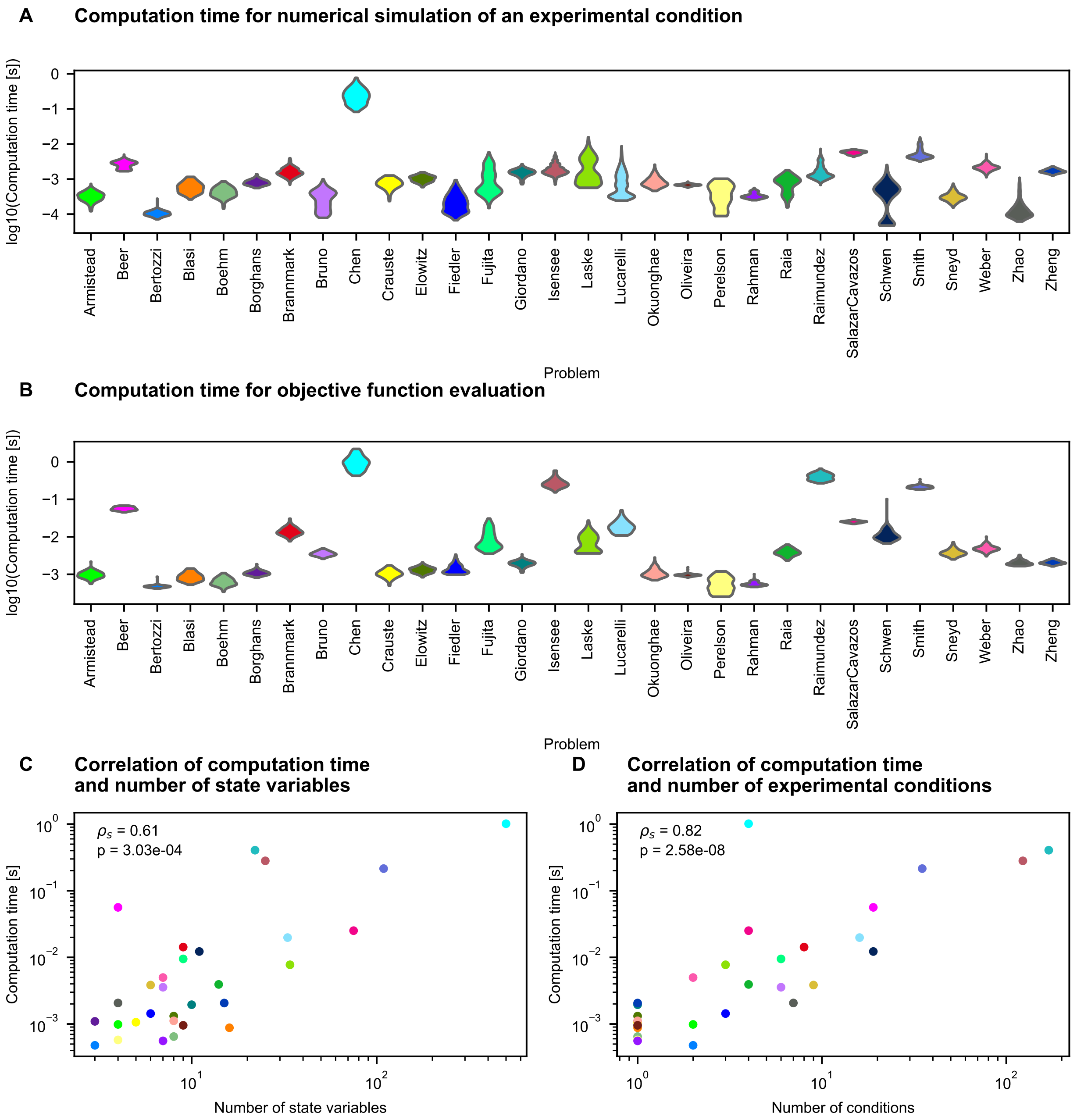

**Supplementary Figure S2. Computation time for simulation and objective function evaluation.** (A) Distribution of computation time required for the numerical simulation of one experimental condition. (B) Distribution of computation time required for the evaluation of the objective function. (C,D) Relation of the average computation time per objective function evaluation and the number of state variables (C) or the number of experimental conditions (D).
To account for the fact that the computation time depends on the parameters, the computations were performed for 100 parameter vectors which are drawn from the prior distribution or uniformly from the parameter space of the respective problem. In (A) and (B) the resulting distributions are shown while in (C) and (D) only average values are shown.

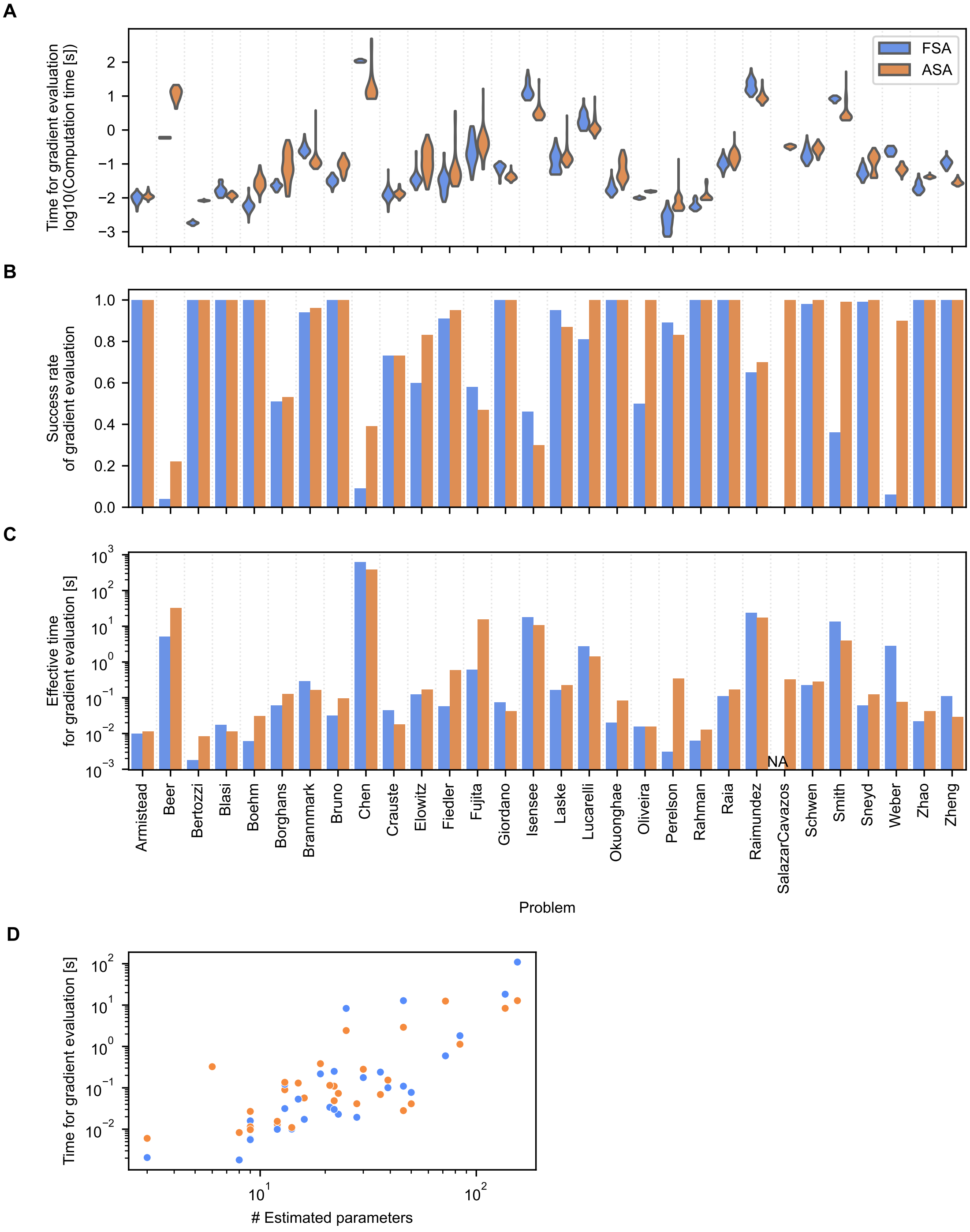

**Supplementary Figure S3. Computation time for objective function gradient evaluation based on forward and adjoint sensitivity analysis.** Results from 100 objective evaluations with parameter vectors sampled uniformly from parameter space. (A) Computation time per objective function gradient evaluation using forward or adjoint sensitivity analysis. (B) Objective function gradient evaluation success rate with forward or adjoint sensitivity analysis. (C) Effective time per objective function gradient evaluation using forward or adjoint sensitivity analysis. The effective time is defined as the total time for the 100 gradient evaluations divided by the number of successful evaluations, and thus, represents the average time required for one successful gradient evaluation. (D) Time per gradient computation over the number of estimated parameters. The median time of the successful objective evaluations is shown.

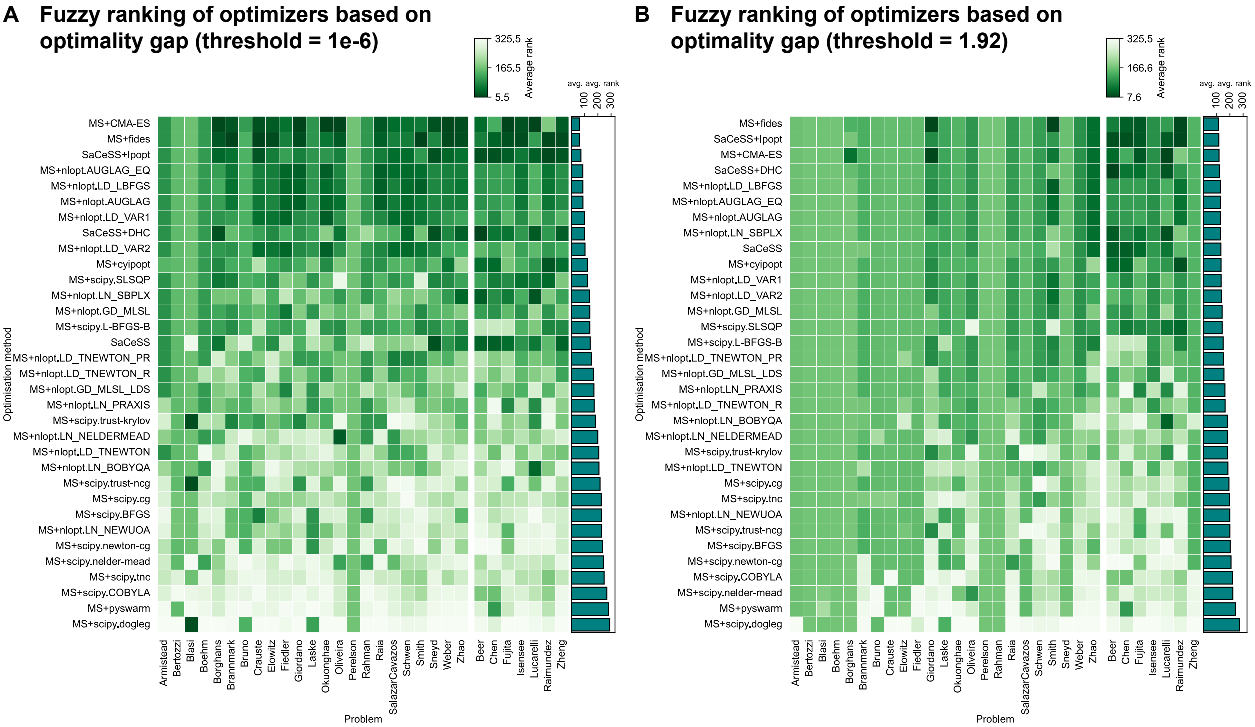

**Supplementary Figure S4. Fuzzy ranking of optimisation methods by optimality gap.** For each problem, we applied fuzzy ranking to the final optimality gaps of all runs and averaged the ranks of the 10 runs per optimisation method. This average rank is shown in the heatmap cells. The bars indicate the column averages. In case of ties in the ranking, the average rank was assigned. Lower values, i.e. darker colours, correspond to better optimisation results. Tolerance for fuzzy ranking: (A) $10^{-6}$ (B) 1.92, corresponding to the $\chi^2$-based rejection threshold at $\alpha = 5\%$.

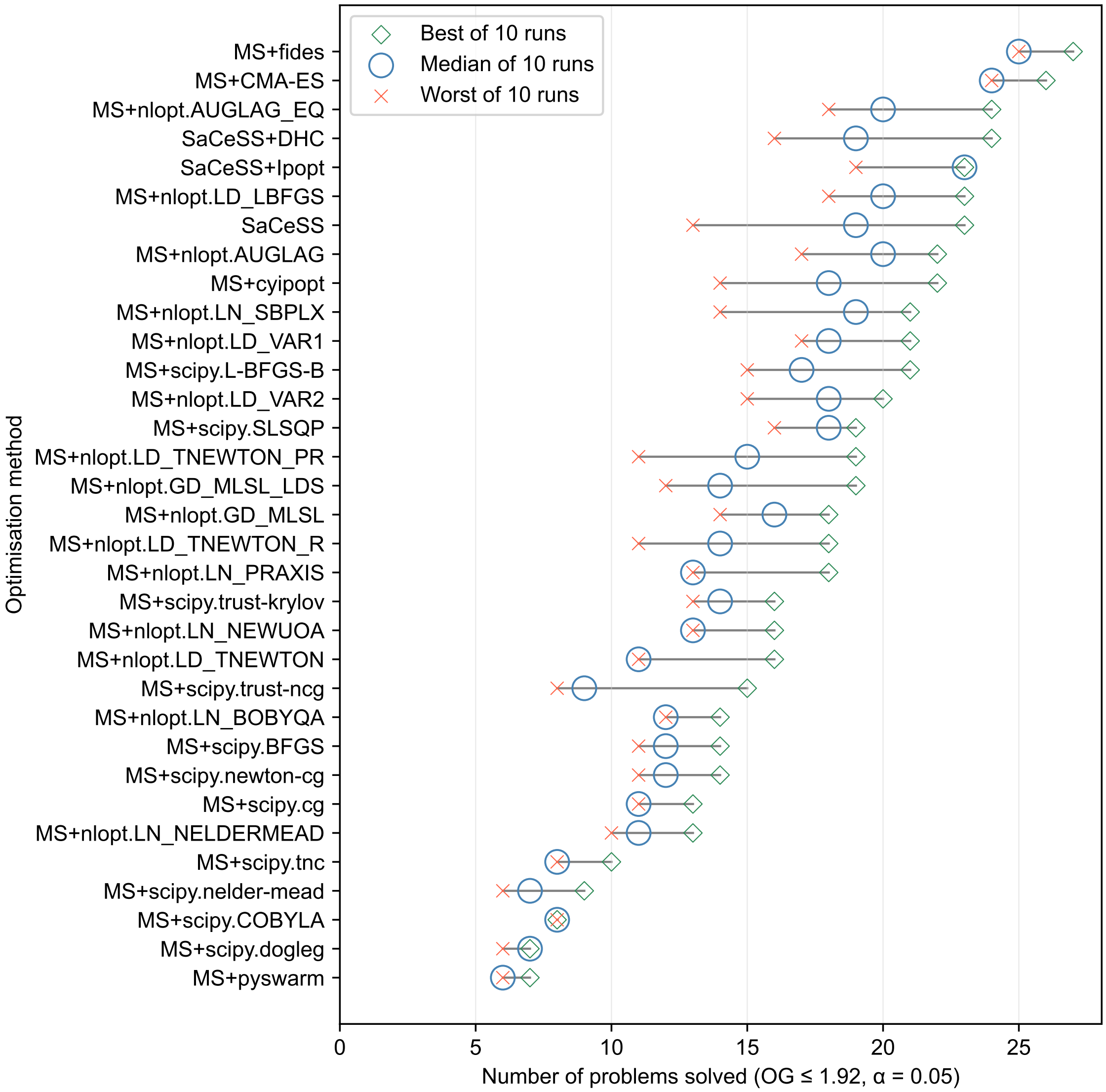

**Supplementary Figure S5. Highest, median, and lowest number of problems solved by optimisation method across 10 runs.** Methods are ranked by the highest, then the median, then the lowest number of problems solved. For each combination of problem and optimisation method, we took the best, median, or worst optimality gap across the 10 runs, and counted a problem as solved if this gap fell below the $\chi^2$-based rejection threshold at $\alpha = 5\%$.

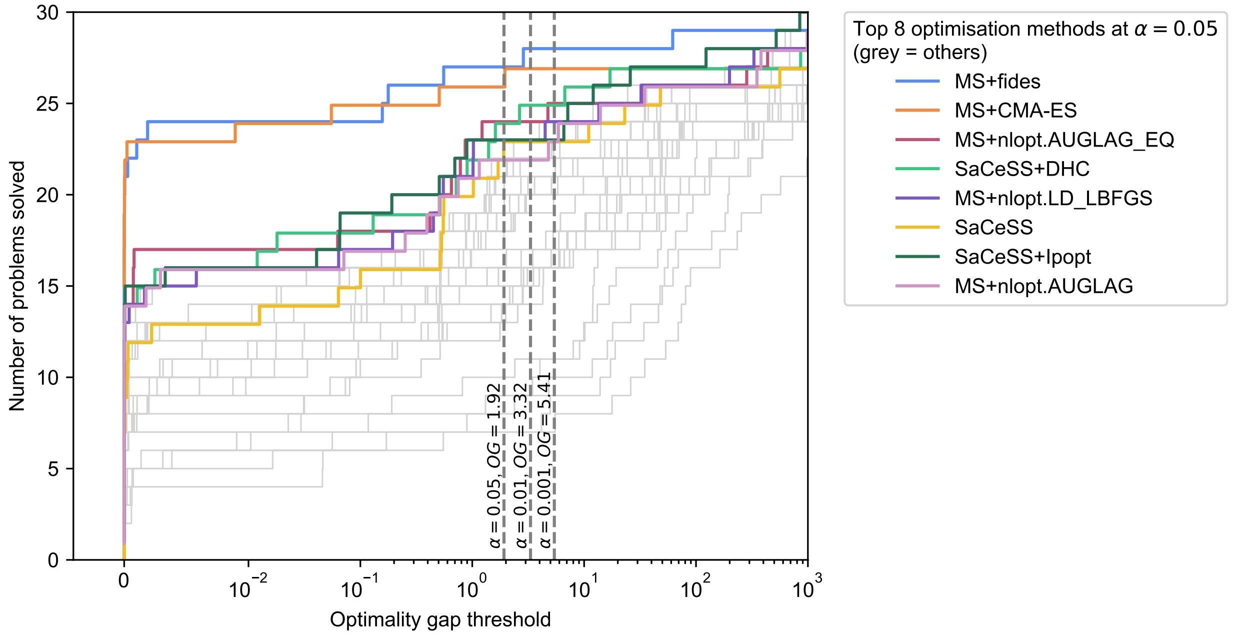

**Supplementary Figure S6. Performance profiles of optimisation methods.** The number of problems solved by each method at various optimality gap thresholds. The dashed vertical lines indicate the optimality gap thresholds for significance levels $\alpha = 0.05$, 0.01, and 0.001. The eight best optimisers at $\alpha = 0.05$ are highlighted, the remaining ones are shown in grey. At each threshold, the minimal optimality gap among the 10 runs is used. The lines are offset slightly to reduce overplotting.

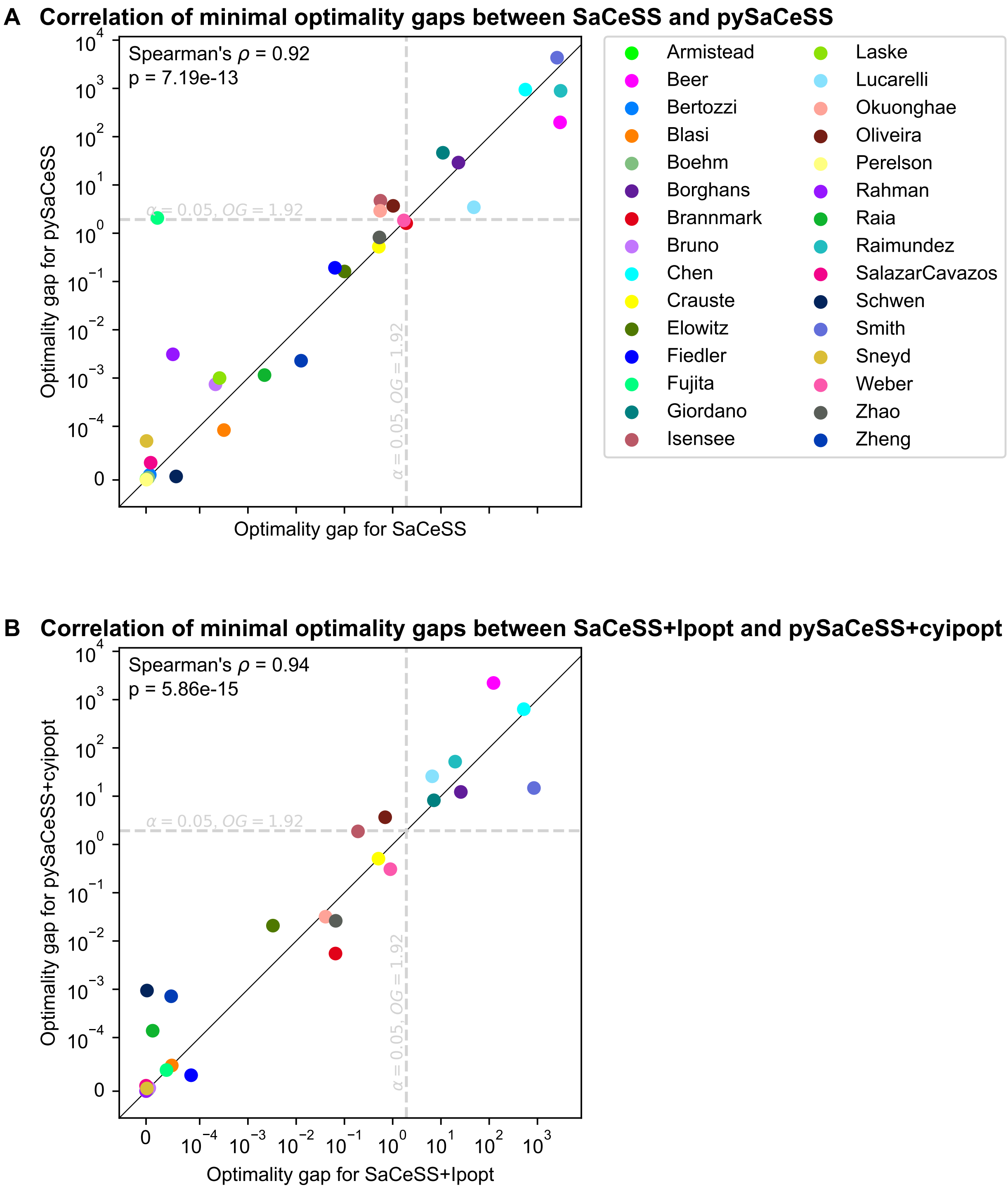

**Supplementary Figure S7. Comparison of optimality gaps of the `SaCeSS` and `pySaCeSS` implementations.** (A) `SaCeSS` without local solver versus `pySaCeSS` without local solver. (B) `SaCeSS` with `Ipopt` as local solver versus `pySaCeSS` with `Ipopt` (through `cyipopt`) as local solver. `Ipopt` was the only optimisation algorithm readily available for use with both the `pypesto` and `parPE` framework. The minimal optimality gaps out of 10 runs are shown. The reference points for the optimality gaps were the best values achieved by any run of the optimisation methods shown in main Figures 1 and 7 on Marvin.

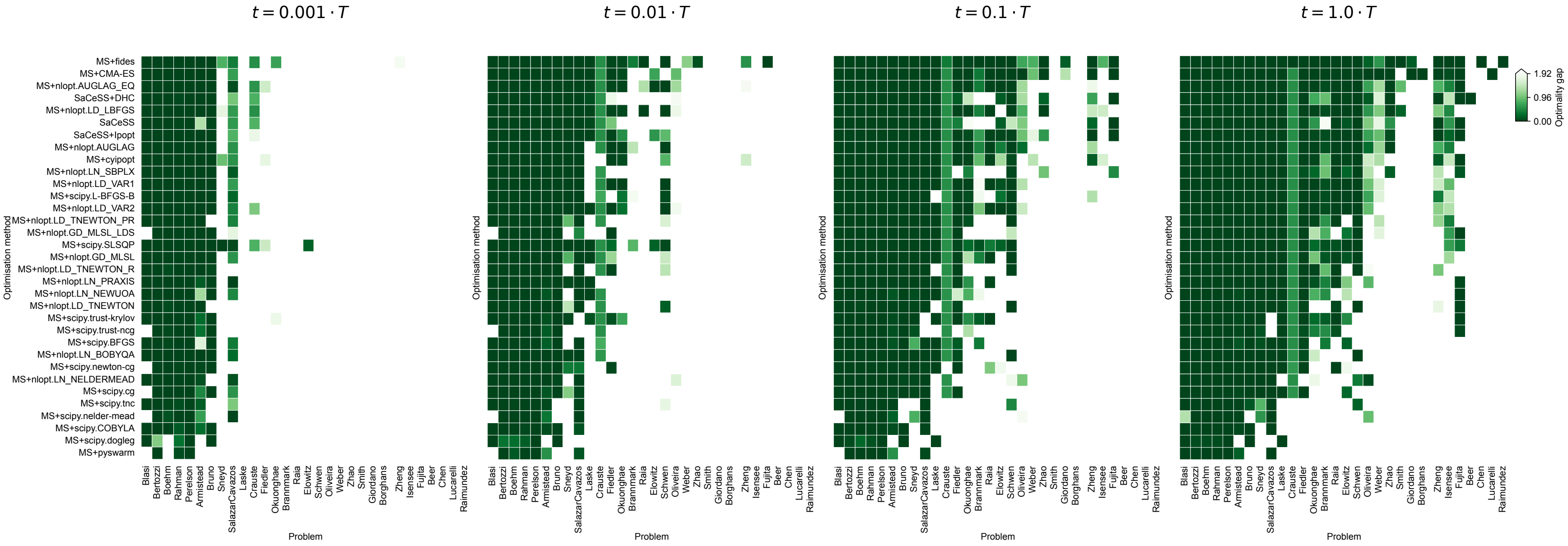

**Supplementary Figure S8. Optimality gaps at different fractions of the time limit.** Heatmaps of optimality gaps for all combinations of problems and optimisation methods based on the best of 10 runs at each timepoint. Greens cells indicate that the problem was solved according to a $\chi^2$ rejection test at $\alpha = 0.05$, white cells indicate the problem was not solved. Optimisation methods are ordered by the number of problems solved within 100% of the compute budget. The row and column ordering is constant across the heatmaps.
